# Rapid establishment of species barriers in plants compared to animals

**DOI:** 10.1101/2023.10.16.562535

**Authors:** François Monnet, Zoé Postel, Pascal Touzet, Christelle Fraïsse, Yves Van de Peer, Xavier Vekemans, Camille Roux

## Abstract

Speciation, the process through which new reproductively isolated species emerge from ancestral populations, occurs due to the gradual accumulation of barriers to gene flow within genomes. To date, the notion that interspecific genetic exchange occurs more frequently between plant species than animals species has gained a strong footing in the scientific discourse. By examining the dynamics of gene flow across a continuum of divergence in both kingdoms, we observe the opposite relationship: plants experience less introgression than animals at the same level of genetic divergence, suggesting that species barriers are established more rapidly in plants. This pattern raises questions about which differences in microevolutionary processes between plants and animals influence the dynamics of reproductive isolation establishment at the macroevolutionary scale.

**One Sentence Summary:** Genetic exchange is more frequent between animal species than plants, challenging historical views.

## Introduction

Introgression, the genetic exchange between populations or between speciating lineages has long been considered an important evolutionary process (*1*). The number of genetic novelties brought by introgression in a population can exceed the contribution of mutation alone, thus increasing both neutral and selected diversity, which can be the source of major evolutionary advances (*2*). One of the consequences of such introgression events is to facilitate the diffusion on a large scale (geographical and/or phylogenetic) of mutations that were originally locally beneficial (*3*). Evidently, genetic exchange across the Tree of Life is not unrestricted but is interrupted by specific mutations, known as species barriers, that reduce hybrid fitness and progressively accumulate in the genome as evolutionary lineages diverge (*4*). These genetic barriers to gene flow contribute to reproductive isolation by either reducing hybrid production or diminishing hybrid fitness. The consequences of reproductive isolation can therefore be captured through the long-term effect of barriers on reducing introgression locally in the genomes, which provides a useful quantitative metric applicable to any organism (*5*). Thus, the genomes of speciating lineages go through a transitional stage, the so-called ‘semi-isolated species’, where they form mosaics of genomic regions more or less linked to barriers to gene flow (*6*). The consideration of this ‘semi-isolated’ status is key to better understanding the dynamics of the speciation process: *i)* When does the transition from populations to semi-isolated species occur? *ii)* At what level of molecular divergence do species become fully isolated?

Historically, reproductive isolation has been studied using complementary approaches. From an “organismal” perspective, it is evaluated using controlled crosses that measure reductions in hybrid viability or fertility, typically over one or two generations (*5*). In contrast, a “genetic” perspective focuses on reductions in gene flow between populations driven by natural selection acting on barrier genes over longer evolutionary timescales (*5*). While laboratory-based approaches offer valuable insights into the genomic architecture of reproductive isolation -such as the number of loci involved, their individual phenotypic effects, and the extent of genetic interactions-they have inherent limitations in detecting species barriers as they occur *in natura* (*7*). To overcome these limitations and investigate the dynamics of reproductive isolation, we examine patterns of gene flow in natural populations, where introgression leaves detectable genomic signatures. Metrics such as *F_ST_* (*8*) and derivatives of the ABBA-BABA test (*9, 10*) are widely employed to assess whether divergence occurred under strict allopatry; however they cannot estimate the timing of gene flow or determine the current isolation status. Recent computational methods explicitly test evolutionary scenarios, including ongoing gene flow and semi-permeable species barriers (*11–13*). Applied to 61 pairs of animal taxa across a continuum of molecular divergence, these methods revealed frequent introgression up to 2% net divergence (*14*) and cases of gene flow between lineages 14 times more divergent than humans and chimpanzees (*15*).

The role of hybridisation and introgression in evolution benefited enormously from the efforts of botanists during the mid-20th century, but the patterns of speciation dynamics described above in animals, specifically the reduction of effective migration between populations or species *in natura*, are still unknown in plants. A historical overview of the literature suggests that plants would be more susceptible to hybridisation, and even introgression, than animals (*2,16–20*). Despite a lack of comparative studies, this notion has been extensively adopted by the scientific community and is supported by some shortcuts. Primarily, the few empirical investigations comparing the dynamics of speciation in plants *versus* animals solely rely on morphological traits to arbitrarily define species (*19*). The emergence of molecular data has now rendered this issue surmountable, as it enables substituting the human-made species concept with genetic clusters that quantitatively vary in their level of genetic distance (*21*) and level of reproductive isolation (*5*). Secondly, the assertion of a higher magnitude of gene flow in plants relative to animals was established without any indication that comparable levels of divergence have been studied. Here, we undertake a comparative genomic approach using molecular datasets from the literature to challenge, with a unified statistical framework, the view that gene flow would be more prevalent in plants than animals for a given level of divergence.

## Results

The present investigation examines the decrease in ongoing gene flow between lineages as a function of their genetic divergence, and compares its dynamics between two kingdoms of the Tree of Life: plants and animals. For this purpose, we empirically explore a continuum of genetic divergence represented by 61 animal pairs and 280 plant pairs. Genomic data from each pair allows the quantification of molecular patterns of polymorphism and divergence by measuring 39 summary statistics commonly used in population genetics and the joint Site Frequency Spectrum (jSFS). For each observed dataset, we then tested whether the observed set of summary statistics was better reproduced by scenarios of speciation with or without migration by using an approximate Bayesian computation framework (ABC; (*11*)). The same ABC methodology for demographic inferences was employed for both the animal dataset (analyzed in (*14*)) and the plant dataset. The new plant dataset was produced from sequencing reads publicly available for 25 genera distributed in the plant phylogeny (212 pairs of eudicots, 45 of monocots, 21 of gymnosperms, 1 lycophyte and 1 magnoliid; Table S1) and were not chosen on the basis of a preconceived idea of their speciation mode (see supplementary materials A.2). The posterior probability of models with ongoing migration computed by the ABC framework is used to assign a status of isolation or migration to each pair along a continuum of divergence (Fig.1-A), allowing the comparison of speciation dynamics between plants and animals. In contrast to the expected outcomes reported in previous studies (*2, 16–20*), our findings suggest that in comparison to animals, plants exhibit a more rapid cessation of genetic exchange at lower levels of genetic divergence. This is characterized by a swifter transition from population pairs that are best-supported by migration models to those that are best described by isolation models (*P <* 1 × 10^−4^; Fig.1-A and table S2). Therefore, by fitting a generalized linear model for the migration/isolation status to the plant and animal datasets, as a function of the net molecular divergence, we determined that at a net divergence of ≈ 0.3% (95% CI: [0.26%-0.36%]), the probability that two plant lineages are connected by gene flow falls below 50%, while in animals this inflection point occurs at higher levels of divergence close to 1.8% (95% CI: [1.1%-2.7%]; Fig.1-A and table S2). The plant dataset comprises genomic data derived from diverse sequencing methodologies, including RAD-sequencing (*n* = 117 pairs), RNA-sequencing (*n* = 111) and whole genome sequencing (*n* = 52), while the animal dataset predominantly consists of RNA-sequencing data (*n* = 52). To control for potential bias in sequencing technologies, we restricted our analysis solely to plant and animal datasets acquired through RNA-sequencing. The key result of a faster cessation of gene flow in plants than in animals is still supported (*P <* 1 × 10^−4^ and table S3), allowing us to reject the idea that our conclusions are derived from such a methodological bias. The number of pairs within a genus showing robust statistical support for either ongoing migration or current isolation in plants ranged from one to 28 pairs. Therefore, we also investigated a possible effect of sampling bias within the plant dataset. Through random sub-sampling involving a single pair of lineages per plant and animal genus, we demonstrate that the contrast in speciation dynamics between plants and animals consistently persists, also rejecting the idea that our result stems from the over-representation of a genus of plants with highly reproductively isolated lineages (Fig. S9).

**Figure 1:**
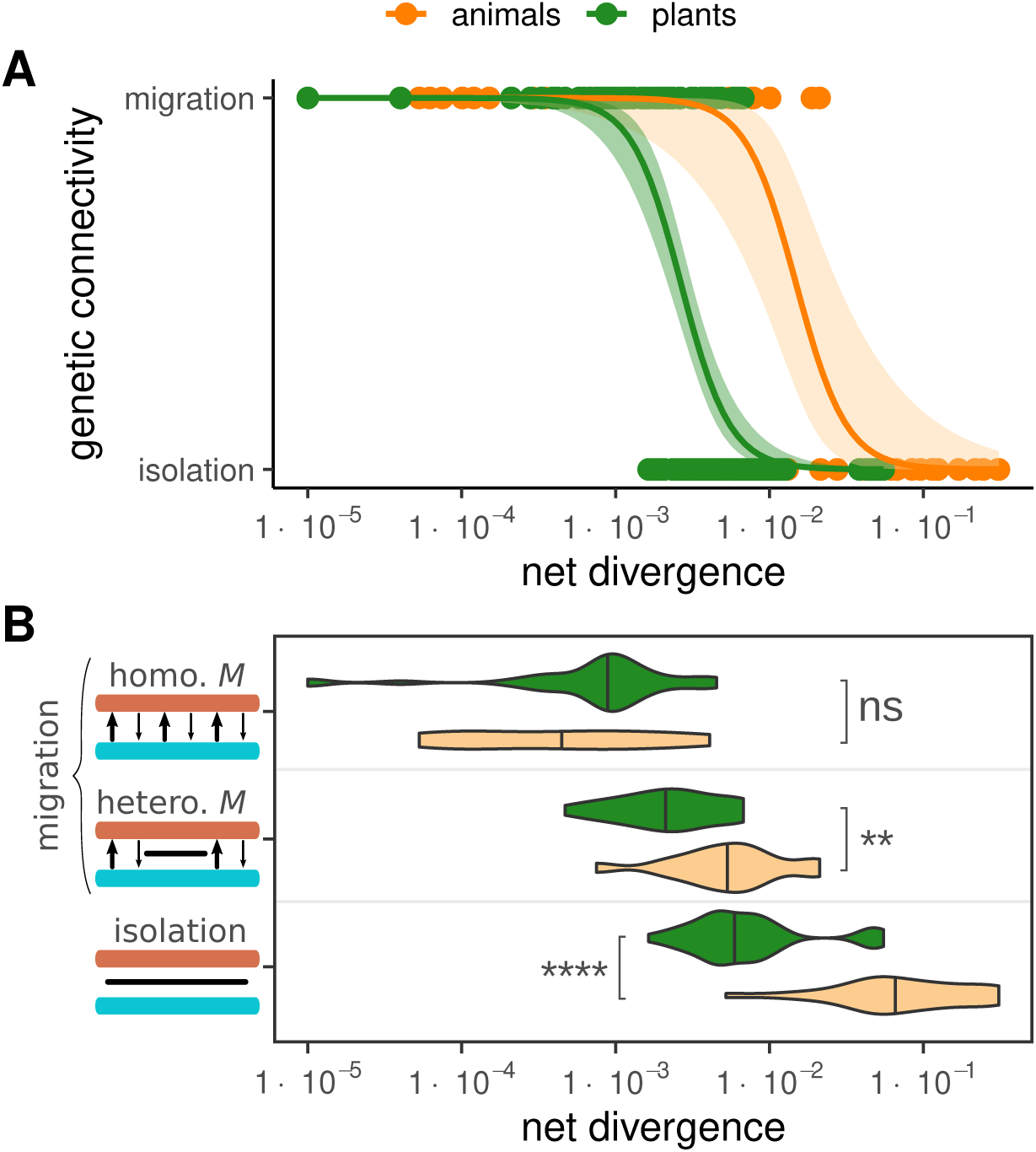
Genomic patterns of introgression along a divergence continuum in plants *versus* animals. Average genomic divergence and migration/isolation status were estimated for 280 plant pairs (green) and 61 animal pairs (orange) using ABC (*14*). **A.** x-axis: net divergence. y-axis: best-supported model (ongoing migration *vs* isolation). Points: population/species pairs. Curves: logit fits for plants and animals. **B.** Net divergence distributions for plant (green) and animal (orange) pairs under homogeneous (homo. M), heterogeneous (hetero. M), or isolation models. Bars along the y-axis: homologous chromosomes (blue, brown), gene flow (arrows), or barriers (black bars). Statistical significance (Mann–Whitney U test): *ns* (*P >* 0.05), * (*P* ≤ 0.05), ** (*P* ≤ 0.01), *** (*P* ≤ 0.001), **** (*P* ≤ 0.0001).

To investigate the build-up of species barriers within the genomes of both plant and animal species, we now focus towards pairs supported by ongoing gene flow. Within the range of speciation scenarios considered, the rate of gene flow can be uniform across genomes (i.e., homogeneous) or it can vary locally from one genomic region to another (i.e., heterogeneous; see Fig. S7), contingent, respectively, upon the absence or presence of barrier genes that are expressed (*6*). The ABC framework described earlier allows us to classify animal and plant pairs as experiencing either genomically homogeneous or heterogeneous introgression (*22*). We find that plants experience a faster shift from the absence of barriers to semi-permeable barriers, the latter occurring at a net divergence of ≈ 0.2% (compared to ≈ 0.6% in animals; Fig.1-B). These findings demonstrate that, in plants, the initial species barriers that generate genomic heterogeneity of introgression rates, as well as the establishment of complete isolation between species, manifest at relatively lower levels of divergence than in animals. This suggests that the speciation process may require fewer mutations in plants than in animals for reproductive isolation to be both initiated and completed, as evidenced by previous experimental crosses unraveling the genomic architecture of reproductive isolation in plants (*23–25*).

Finally, we conducted a comparative analysis of the temporal patterns of gene flow during divergence in plants *vs* animals. We specifically examine whether ongoing gene flow predominantly arises from a continuous migration model, initiated since the subdivision of the ancestral population (as illustrated in Fig. 2), or if it is a consequence of secondary contact following an initial period of geographic isolation and divergence. This model comparison using ABC is restricted to pairs for which we previously found a strong statistical support for ongoing gene flow. Our analysis shows that plants and animals differ in their primary mode of historical divergence, specifically in the extent of gene flow during the initial generations after the lineage split. Indeed, among animals, roughly 80% of the pairs that exhibit robust statistical evidence of ongoing migration diverged in the face of continuous gene flow since the initial split from their ancestor (Fig. 2). A minority of animal pairs (20%) underwent primary divergence in allopatry before coming into secondary contact. Conversely, in the case of plants, pairs that display ongoing gene flow have more frequently experienced secondary contacts (≈ 52%; Fig. 2), in line with what is commonly assumed in plants (*26*). To ensure that the observed pattern of reduced gene flow in plants compared to animals is not merely a consequence of greater geographic distances between plant pairs, we calculated the minimum geographic distance between taxa within each pair using the GPS coordinates of the sampled individuals (Fig. S1). Strikingly, our analyses reveal that ongoing migration is less frequent in pairs of plant lineages despite their closer average minimum geographic distance (≈ 488 km) than in animals (≈ 2, 230 km), confirming that current geography is a poor predictor of genetic introgression in the history of sister species both in plants (*P* = 0.936) and animals (*P* = 0.688).

**Figure 2:**
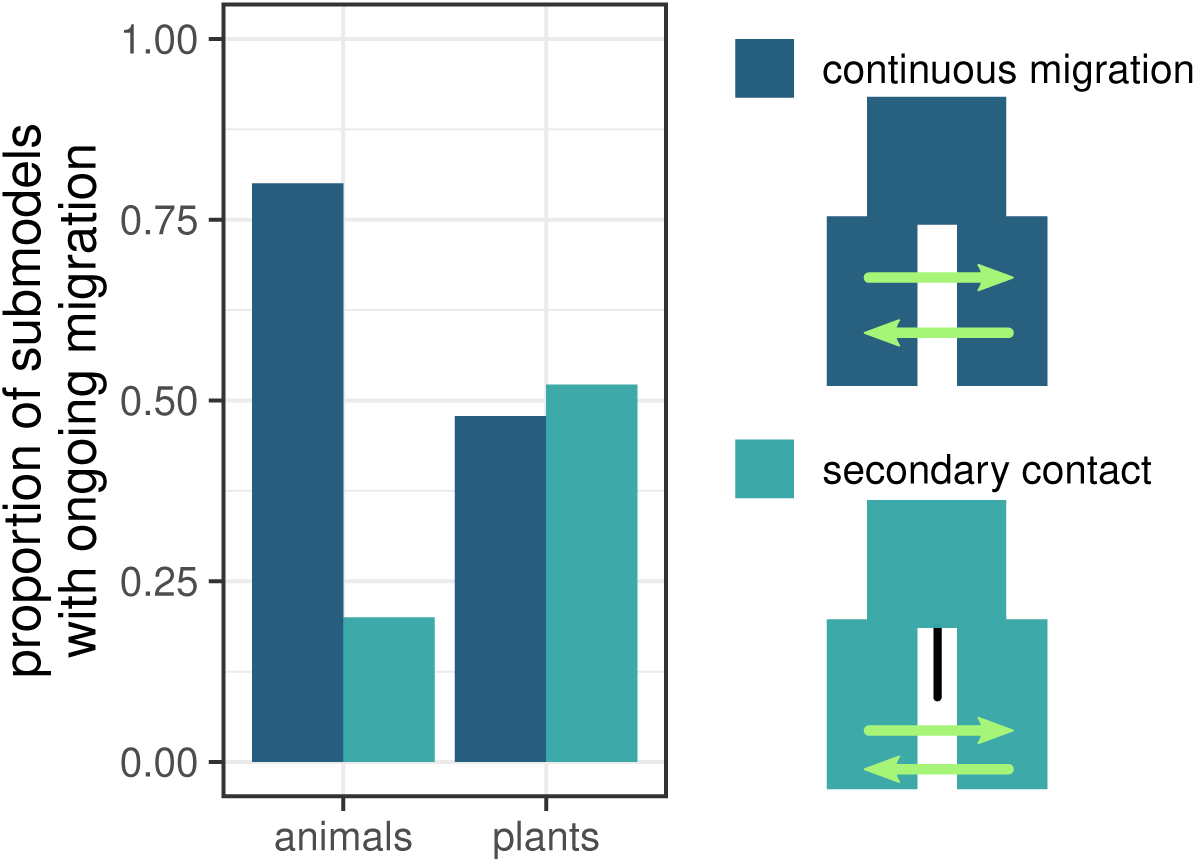
Temporal models of ongoing migration in plants and animals. Strong statistical support of ongoing migration was observed in 69 pairs of plants and 30 pairs of animals. For every pair supported by ongoing gene flow, sub-models were compared to distinguish between continuous migration (dark blue) *versus* secondary contact (light blue).

## Discussion

The historical literature on hybridisation defined hybrids as the offspring of crosses between individuals from genetic lineages “*which are distinguishable on the basis of one or more heritable characters*” (*27*). Within this conceptual framework, examinations of numerous wild species have demonstrated a greater incidence of interspecific hybridization in plants than in animals (*19*), thus supporting the original assumption that plants are indeed more likely to hybridize than animals (*2, 16*). However, the advent of molecular markers to measure genetic differentiation in the early 2000s provided results in contrast to morphological studies, particularly by illuminating the higher *F*_ST_ values within plant species relative to animals (*28, 29*), indicating higher gene flow at the intraspecific level in the latter. Moreover, Morjan and Rieseberg (*28*) showed that this difference between kingdoms persists regardless of the mating system (from outcrossing to selfing) or the geographical distribution (local, regional, or biregional ranges). In our methodological approach, we depart from the human-made conception of ‘species’ and instead focus on genetic clusters that exhibit varying degrees of divergence and varying degrees of connectivity due to gene flow (Fig. S4). We could only attain this level of resolution because our methodology explicitly models the divergence history between lineages and captures the effect of species barriers on genomic patterns of gene flow. In doing so, we unravel the apparent paradox between studies of reproductive isolation between morphologically differentiated entities that suggest more frequent hybridization events in plants than in animals, and the greater genetic differentiation observed within plant species with molecular markers. Indeed, our explicit comparisons of ongoing migration models support the idea that scenarios of secondary contact are particularly frequent among the surveyed lineages in plants (*26*), whereas pairs of closely related animal species tend to experience gene flow more continuously over time. Secondary contact scenarios involve a preliminary phase of allopatry which affects the divergence of sister lineages across all markers: molecular and morphological. Such a historical context may thus engender the misconception that plants undergo hybridization events more frequently than animals, simply because these events are more conspicuous in plants, as introgression happens more often between morphologically distinct lineages experiencing secondary contact. This result suggests, conversely, that genetic introgression in animals may often be overlooked, particularly if it occurs more frequently between species lacking clear morphological differences compared to plants. However, we strongly emphasize that our findings should not be interpreted as evidence that speciation in animals frequently occurs in the presence of gene flow. The gene flow inferred here is a proxy for reproductive isolation, reflecting barriers that, on average, accumulate more slowly across animal genomes.

The speciation process clearly does not follow a universal molecular clock, although certain molecular constraints inevitably make the process irreversible once a certain level of divergence is reached (*4, 19*). While light has recently been shed on the rarity of hybrid zones found in plants (*26*), another mystery has now been added: why is the probability of being reproductively isolated greater in plants than in animals given identical genomic divergence? The speciation process is inherently multifactorial, involving a combination of factors such as ecological, genetic, geographical, and cytological, among others. Our study relies exclusively on a comparative population genomics approach, which is insufficient to provide an integrative explanation for why plants appear to accumulate reproductive isolation more rapidly than animals. Addressing this complex question will require future studies that combine diverse approaches across multiple models (*30, 31*). Here, we propose non-exclusive and non-exhaustive hypotheses to explain why reproductive isolation appears to accumulate faster in plants than in animals, with further details provided in the supplementary material (section B):

1. **Chloroplast as an additional source of barriers:** The plant-specific chloroplast organelle drives species-specific co-evolution with the nuclear genome, accumulating compensatory mutations necessary for nucleo-cytoplasmic interactions (*32*). In hybrids, mismatched nuclear and chloroplast genomes can cause developmental disruptions that reduce gene flow between closely related plant species (*33*).
2. **Mitochondrial mutation rates:** Mitochondrial mutation rates are generally higher in animals than in plants (*34*). However, higher mutation rates tend to relax compensatory selection on nuclear mutations. As a result, plants may accumulate more mito-nuclear incompatibilities than animals, contributing to their reproductive isolation (*35*).
3. **Selfing rates and reproductive isolation:** Plants display diverse mating systems, from high selfing to high outcrossing (*36*). Selfing species exhibit greater speciation rates (*37*), likely due to reduced gene flow, smaller effective population sizes, and higher linkage disequilibrium (*38*). These factors may facilitate faster reproductive isolation in selfing plants compared to outcrossers (*39*). In our dataset, we do not observe a strong effect of the reproductive system on the dynamics of reproductive isolation establishment (Fig. S13; Table S4), primarily due to the bias toward outcrossers, which does not adequately reflect the variance in selfing rates among plants (Fig. S12).
4. **Pollinator shifts and reproductive isolation:** Shifts in pollinator preferences, particularly among allopatric plant lineages, can drive pre-zygotic isolation and reduced hybridization rates (*40*). This is reflected in higher diversification rates in plants with fewer specialized pollinators (*41*).
5. **Dispersal modalities:** Plant dispersal through passive vectors (e.g., pollen and seeds) leads to higher genetic differentiation among populations compared to the active mobility of many animals (*28*). This differentiation can promote partial reproductive isolation through outbreeding depression (*42*).
6. **Haploid selection:** Plants may experience stronger haploid selection during pollen development compared to the limited haploid selection in animals (*43*). This difference could amplify the effects of recessive reproductive isolation barriers in plants (*44, 45*).
7. **Parental conflicts during seed development:** Polyandrous reproduction in plants creates intragenomic conflicts during seed development, leading to co-evolution of selfish elements and their suppressors (*46*). These conflicts can create severe hybrid seed development issues, contributing to species barriers (*47*).
8. **Genetic architecture of reproductive isolation:** Quantitative genetic studies reveal that reproductive isolation in plants often involves simpler genetic interactions compared to animals, with fewer loci exhibiting strong individual effects (*23–25, 48*). Additionally, alleles associated with reproductive isolation in plants frequently exhibit intraspecific polymorphism, indicating that stochastic processes often maintain these alleles in populations until their fixation or loss. In contrast, speciation genes in animals are more likely to show signatures of positive selection, suggesting stricter conditions for the establishment of barriers in animals (*49*).

The proposed factors presented here are evidently not mutually exclusive, and it would be misleading to attribute the differences in speciation dynamics between plants and animals to a single, easily testable factor. Understanding which properties of plants and animals, acting at the micro-evolutionary scale, drive such disparities in speciation patterns at the macro-evolutionary scale remains a fundamental challenge that calls for a long-term, community-based initiative for integrative speciation research across fields and taxa (*31*).

A critical question emerging from our findings is whether the accelerated accumulation of reproductive barriers at the population level—leading to more rapid reductions in gene flow in plants compared to animals—could be a key driver of the higher average speciation rates observed in plants at phylogenetic scales (*50*). Addressing this question would bridge the micro- and macro-evolutionary perspectives, offering new insights into how differences in reproductive isolation dynamics shape the tempo and mode of diversification across taxa.

Finally, the methodology employed in this study to investigate variations in speciation dynamics between plants and animals provides a versatile framework that can be extended to examine other contrasts involving diverse life-history traits, such as external *versus* internal fertilization, self-fertilization *versus* cross-fertilization, or haplobiontic *versus* diplobiontic life cycles. Future studies leveraging this framework would help disentangle the roles of these biological factors in the establishment and maintenance of reproductive isolation, further advancing our understanding of speciation processes.

## Acknowledgments

We are grateful to the Institut Français de Bioinformatique (IFB; ANR-11-INBS-0013) thanks to whom bioinformatics analyses and demographic inferences have been carried out.

The authors would like to thank Hélène Morlon, Bert Van Boxlaer, John Pannell and Martin Lascoux for their constructive feedback.

## Authors contributions

F.M (data curation, formal analysis, investigation, visualization, writing), Z.P (data curation), P.T (resources, data curation, supervision), C.F (methodology, writing), Y.V.d.P (funding acquisition, supervision, writing), X.V (funding acquisition, supervision, writing), C.R (funding acquisition, conceptualization, methodology, supervision, coding, investigation, visualization, writing).

## Data and materials availability

All data are available in the manuscript or the supplementary materials. The assembled datasets, the list of references used for mapping and the results of demographic inference are deposited in `Zenodo` with the DOI (https://doi.org/10.5281/zenodo.14365924) (*51*). Scripts for bioinformatic treatment of raw sequencing data are available from GitHub https://github.com/Ladarwall/Greenworld.

## Supplementary Materials

### A Materials and Methods

#### A.1 Animal dataset

The animal data come from the Roux et *al.* (2016) study (*14*). They consist essentially of non-model animal populations/species, initially selected without any particular knowledge about the demographic history, and were sampled from natural populations. These data were produced by RNA sequencing, and only synonymous positions were retained for statistical inferences.

#### A.2 Plant dataset

Raw data used in this work comes from previously published studies (*52–77*). The following criteria were applied to identify datasets in plants:

1. Currently diploid genomes.
2. High-throughput sequencing, i.e, RNA-seq, RAD-seq or whole genome sequencing (WGS).
3. Freely available from `NCBI`.
4. Individuals sampled from natural populations (geographic distribution represented in Fig. S1).
5. A minimum of two sampled populations/species per genus.
6. A minimum of two sequenced individuals per sampled population/species.

Datasets fitting these criteria were examined through exploration of literature found *via* Google Scholar (https://scholar.google.com), `NCBI` (https://www.ncbi.nlm.nih.gov/Traces/study/) and `DDBJ` (https://ddbj.nig.ac.jp/search).

Finally, 118 different plant species/populations from 25 different genera were retained for the demographic analysis according to our criteria (Table S1), allowing 280 pairwise demographic analyses to be carried out. These comparisons cover all possible pairs within each genus. No comparisons are made between different genera, with the exception of comparisons within the *Laccospadicinae* (*Howea* and *Linospadix*) due to their relatively small genetic distance.

#### A.3 Assembly, read mapping and genotype calling

For the plant datasets: reads and metadata were downloaded using SRA-Toolkit, version 2.11.0 (https://github.com/ncbi/sra-tools/wiki/01.-Downloading-SRA-Toolkit). Here we separate plant projects for which we worked with synonymous positions (from RNA-seq: *n*=7 genera and WGS: *n*=4) from those for which we could not (from RAD sequencing: *n*=13):

##### A.3.1 Reads from RNA-seq and WGS

In line with the animal dataset (*14*), the bioinformatic strategy applied to the plant data is to retain synonymous positions. Reads for a given population/species pair were therefore mapped to a reference transcriptome with the bowtie2 program version 2.4.2 (*78*): either taken from the `1KP project` (*79*) if a species of the same genus is represented there (https://db.cngb.org/onekp/search/), or taken from the data associated with the original articles when available (Table SS1). Every position (variants and invariants) were called with a minimum of 8 reads using Reads2SNP 2.0, the uncalled low-quality positions were then coded as “N”. The resulting fasta file was used for each population/species as the input file for the demographic inferences.

##### A.3.2 Reads from RAD-seq

Loci were assembled for each RAD-seq dataset using `Stacks` 2.6 (*80, 81*). Combinations of parameters were explored following Paris et *al.* 2017 (*82*) to maximise the amount of biological information retained. Using the two or four samples with the highest amount of available data per lineage, assemblies were built using *denovo map.pl* (`Stacks`) with different combinations of parameters: the minimum depth for a stack to be valid (*-m*, ranging from 3 to 5), the number of mismatches allowed between stacks within individuals (*-M*, ranging from 1 to 6) and the number of mismatches allowed between stacks between individuals (*-n*, set to *M* or *M* + 1), for a total of 36 combinations. In addition, loci that were missing in at least 20% of the samples per population were withdrawn with the argument *–min-samples-per-pop 0.80* (i.e. only loci with the information for all samples were kept, as populations were composed of two or four samples). The number of polymorphic loci was plotted as a function of the different combinations of parameters using a homemade `R` script. For each dataset, a combination was selected in function of the trade-off between maximising the number of polymorphic loci and minimising the parameter values to produce a reference set of loci for each species/population pair. Reads were mapped on this reference with bowtie2 version 2.5.1, and variants were called with Reads2SNP 2.0 in the same way as “RNA-seq and WGS” datasets.

#### A.4 Demographic inferences

##### A.4.1 Choice of the inferential method

Several methods exist to test introgression between evolutionary lineages, which can broadly be categorized into population genomic approaches (e.g., FastSimCoal (*83*), *∂a∂i* (*84*), Moments (*85*), DILS (*11*)) and phylogenetic methods (e.g., QuIBL (*86*), Dsuite (*87*)). All these methods are effective within the contexts for which they were designed. The goal of this section is not to demonstrate the superiority of one approach over another but to justify why DILS was chosen as a relevant tool for addressing the specific question posed in this study: whether there is ongoing gene flow between two populations or whether they are currently isolated. The choice of method also depends on the properties of the sampling (i.e., number of sampled individuals, number of sampled populations in the genus, assumptions about demography for certain population pairs, etc.) as well as on the properties of the molecular data (i.e., locus length, presence or absence of intra-locus recombination, etc.). Finally, we also aimed to reduce biases that arise when linked selection is not taken into account, whether from background selection (*88*) or selection against species barriers (*89*). In the next section, we focus on a comparison between QuIBL and DILS because the literature has shown that FastSimCoal and *∂a∂i* produce results largely comparable to DILS when applied to the same datasets (*90, 91*).

###### Key methodological differences between DILS and QuIBL

In a given phylogenetic tree with more than three lineages (species or populations), QuIBL extracts information from individual gene trees and tests, for a given triplet of lineages, whether the distribution of internal branch lengths can be explained solely by incomplete lineage sorting or whether introgression processes must also be invoked. For a tree of the topology ((1, 2), 3), QuIBL is particularly powerful in detecting deviations from strict allopatry between lineages 1 (or 2) and 3. Such deviations are interpreted as evidence of historical introgression events, which may vary in recency.

In contrast, DILS leverages population genomic summary statistics, particularly those derived from the Site Frequency Spectrum (SFS), to evaluate whether observed patterns are better explained by a model with ongoing migration or one without, using an Approximate Bayesian Computation (ABC) framework. Given that the question addressed here concerns the extent of current reproductive isolation between specific pairs of lineages (in plants and animals), it was important to discriminate between different temporal patterns of introgression. Deviations from strict allopatry caused by ancient migration do not convey the same implications for current reproductive isolation as those caused by secondary contact. This distinction is explicitly addressed by DILS through the analysis of intra-specific polymorphism and interspecific divergence data for a given pair of lineages.

###### Rationale for Choosing DILS

Our decision to use DILS was motivated by the following considerations:

- DILS accounts for variations in effective population size over time, whereas phylogenetic approaches assume a constant *N_e_*.
- DILS accounts for genome-wide variations in effective population size caused by background selection, as well as variations in effective migration rates driven by linked selection against species barriers.
- DILS accounts for intra-locus recombination, while the distribution of internal branch lengths is derived under the assumption of a simple tree per locus, with no recombination occurring within individual loci.
- DILS does not require a specific sampling scheme to test introgression, whereas phylogenetic methods cannot detect gene flow between lineages 1 and 2 in a tree of the topology ((1, 2), 3).
- DILS works directly with measurable quantities derived from sequences (e.g., *F_ST_*, *θ_W_*, *π*, *D_a_*, *D_xy_*, etc.) without requiring a preliminary step of inferring phylogenetic trees for each locus.
- DILS can handle multiple individuals within each species, reducing the risk of overlooking shared polymorphisms between species.
- DILS does not require long loci for accurate inference, making it less sensitive to biases introduced by RADseq or RNAseq data. In contrast, phylogenetic methods may perform poorly with short loci and are subject to violations of no-recombination assumptions with long loci (*92*). In addition, very long loci may buffer local genomic variations for *N_e_* and *N_e_.m*.
- DILS works directly on pairs of lineages and does not require data from more than three species, allowing its application to genera where only two species have been sequenced.

###### Comparison between DILS and QuIBL on simulated data

To further illustrate the relative strengths of these approaches, we conducted a comparative analysis of DILS and QuIBL using coalescent simulations under a four populations model (Fig. S5). Data were simulated under various temporal patterns of introgression using `msnsam` (*93*), a modified version of `ms` (*94*). The simulations were designed to satisfy all assumptions made by QuIBL (e.g., long loci, no intra-locus recombination, no migration between lineages 1 and 2 in a tree of the topology ((1, 2), 3)). For QuIBL, the input consisted of exact, simulated gene trees rather than inferred trees, as would be the case with real biological data.

###### Simulated Scenarios

Four demographic scenarios were simulated (Fig. S5):

- **Strict Isolation (SI_4pop_)**: No migration.
- **Isolation-Migration (IM_4pop_)**: Migration between lineages 2 and 3 from the present to *T*_4_.
- **Ancient Migration (AM_4pop_)**: Migration between lineages 2 and 3 between *T*_dem_ and *T*_4_.
- **Secondary Contact (SC_4pop_)**: Migration between lineages 2 and 3 from the present to *T*_dem_.

Migration was modeled symmetrically at a rate *M* = *N_e_.m*, where *N_e_* is the effective population size and *m* is the proportion of migrants in each generation.

The parameters explored included:

- *T*_4_: Speciation time between lineages 1 and 2 (0.5, 2, 4 *N_e_* generations).
- *T*_3_: Speciation time between (1, 2) and 3 (*T*_4_ + 0.5, 2, 4 *N_e_* generations).
- *T*_2_: Speciation time between ((1, 2), 3) and 4 (40 *N_e_* generations).
- *T*_dem_: Migration onset/end time (*T*_4_ · [0.1, 0.25, 0.5] for SC_4pop_; *T*_4_ · [0.5, 0.75, 0.9] for AM_4pop_).
- *M* (= *N_e_.m*): Number of migrants per generation (0.25, 10 migrants).
- *L*_mig_: Number of loci affected by migration (100, 500, 1000 loci out of 1,000 total loci).

Migration is assumed to be symmetric, occurring at a rate *M* corresponding to *N_e_.m*, where *N_e_* is the effective population size and *m* is the proportion of individuals in a population consisting of migrants in each generation. We simulate 1,000 independent loci, with no intra-locus recombination. Among these loci, a subset *L*_mig_ is influenced by migration, while the remaining loci represent genetic isolation between the species.

It is important to note that the assumptions of a constant *N_e_* over time and across the genome, as well as the absence of intra-locus recombination, are not required by DILS. These constraints are therefore relaxed in the empirical application to plants *versus* animals. However, they are implemented in the simulation study to enable a direct comparison between DILS and QuIBL, as they align with the assumptions required by QuIBL.

Simulations for QuIBL and DILS were performed using the same parameter combinations. Differences lay in sampling schemes: for QuIBL, a single copy per locus was sampled from lineages 1, 2, 3, and 4, with exact gene trees simulated and rooted using lineage 4 (Fig. S5). For DILS, two diploids were sampled from lineages 2 and 3, reflecting the minimal sampling scheme when DILS was applied to the empirical plants *versus* animals dataset.

Simulations were generated using the script `simulations_for_QuIBL_DILS_full. py`. The command lines for simulating data using `msnsam` under different demographic models are as follows:

**SI**_4*pop*_:

- **QuIBL:** `msnsam 4 1000 -T -I 4 1 1 1 1 0 -ej tbs 2 1 -ej tbs 3 1 -ej 10 4 1`
- **DILS:** `msnsam 8 1000 -t 40 -I 4 0 4 4 0 0 -ej tbs 2 1 -ej tbs 3 1 -ej 10 4 1`

**AM**_4*pop*_:

- **QuIBL:** `msnsam 4 1000 -T -I 4 1 1 1 1 0 -em tbs 2 3 tbs -em tbs 3 2 tbs -eM tbs 0 -ej tbs 2 1 -ej tbs 3 1 -ej 10 4 1`
- **DILS:** `msnsam 8 1000 -t 40 -I 4 0 4 4 0 0 -em tbs 2 3 tbs -em tbs 3 2 tbs -eM tbs 0 -ej tbs 2 1 -ej tbs 3 1 -ej 10 4 1`

**IM**_4*pop*_:

- **QuIBL:** `msnsam 4 1000 -T -I 4 1 1 1 1 0 -m 2 3 tbs -m 3 2 tbs -eM tbs 0 -ej tbs 2 1 -ej tbs 3 1 -ej 10 4 1`
- **DILS:** `msnsam 8 1000 -t 40 -I 4 0 4 4 0 0 -m 2 3 tbs -m 3 2 tbs -eM tbs 0 -ej tbs 2 1 -ej tbs 3 1 -ej 10 4 1`

**SC**_4*pop*_:

- **QuIBL:** `msnsam 4 1000 -T -I 4 1 1 1 1 0 -m 2 3 tbs -m 3 2 tbs -eM tbs 0 -ej tbs 2 1 -ej tbs 3 1 -ej 10 4 1`
- **DILS:** `msnsam 8 1000 -t 40 -I 4 0 4 4 0 0 -m 2 3 tbs -m 3 2 tbs -eM tbs 0 -ej tbs 2 1 -ej tbs 3 1 -ej 10 4 1`

The QuIBL analyses were performed using the following options:

- `-numdistributions: 2`
- `-likelihoodthresh: 0.01`
- `-numsteps: 10`
- `-gradascentscalar: 0.5`
- `-multiproc: False`
- `-maxcores: 1000`
- `-totaloutgroup: 4`

For the DILS inferences on the simulated datasets, we used the following parameters: assuming a constant population size (`population growth: constant`), a mutation rate *µ* of 2 × 10^−8^ mutations per generation per nucleotide (`mu: 0.00000002`), no intra-locus recombination (`rho over theta: 0`), and an effective population size *N_e_* uniformly explored between 0 and 200,000 individuals. The split time was uniformly explored between 0 and 1,500,000 generations, and migration (*N_e_.m*) was uniformly explored between 0.2 and 10 migrants per generation. No outgroup was used to polarize mutations (`nameOutgroup: NA`).

The final output file, table QuIBL DILS.txt, summarizes the demographic parameters and the inference results obtained from DILS and QuIBL for each simulated dataset.

###### Results of inferences performed with DILS and QuIBL on simulated datasets

When the simulated model represents current isolation (SI_4_*_pop_* and AM_4_*_pop_*), both methods converge on the correct model in approximately 75% of simulations across all explored parameter combinations (Fig. S6-A, left panel). DILS performs slightly better than QuIBL, with around 14% of datasets where it is the only method to correctly recover the isolation model, compared to approximately 6.5% for QuIBL. However, these performances in supporting isolation are of the same general magnitude for both methods. For the parameter combinations used in these simulations, both methods fail to recover the isolation model in about 4.5% of cases.

The differences between the two methods become more pronounced when the true model involves ongoing migration (IM_4_*_pop_* and SC_4_*_pop_*). In this scenario, both methods converge on the correct model in approximately 36% of simulations under the explored parameter combinations (Fig. S6-A, right panel). QuIBL alone identifies migration in only 0.001% of simulations, while DILS is the sole method to correctly infer migration in about 29% of cases. This analysis suggests that both methods are highly conservative, failing to detect migration in 35% of simulations with ongoing gene flow. However, it is important to note that these results stem from discrete parameter combinations (*T*_4_, *T*_3_, *T*_dem_, *M*, and *L*_mig_), where migration can be both low (*N_e_.m* = 0.1) and affecting only a small portion of the genome (10% of loci).

When we focus on simulations with higher levels of introgression (Fig. S6-B, right panel), the performance of both methods improves significantly. In these cases, the two methods converge on the correct model with ongoing migration in approximately 72% of simulations. The remaining simulations with ongoing migration are correctly recovered solely by DILS, and no simulations are simultaneously missed by both methods.

Our choice to use DILS in this study was primarily driven by its demonstrated reliability in previous analyses and its ease of use for interpreting temporal patterns of introgression via explicit scenario testing. Unlike QuIBL, which excels in handling large phylogenies, DILS is particularly well-suited for datasets with simpler sampling schemes, such as cases where only two species are available for analysis. Our comparison confirms that our initial choice of DILS did not come at the expense of reduced performance. This validation, combined with DILS’s interpretability and suitability for the scope of our study, highlights its relevance for addressing the research questions we aimed to explore.

##### A.4.2 DILS

Model comparisons were carried out using the approximate Bayesian computation (ABC) framework applied in the animal study (*14*) and distributed under the name DILS (Demographic Inferences with Linked Selection (*11*)). DILS aims to test whether the studied species are connected by gene flow. It incorporates the effects of linked selection by modeling background selection as heterogeneous effective population sizes (hetero-*N_e_*) across the genome and selection associated with barriers to gene flow as genomic heterogeneity in effective introgression rates (hetero-*N_e_.m*). Here we describe how DILS works.

##### A.4.3 Compared models

The primary objective of our demographic analysis is to determine which two populations historical scenario explain the best a given dataset. The term dataset here refers to a pair of populations/species (comprising either two animal or two plant lineages), for which genomic data are described by an array of summary statistics (see section A.4.4). In our ABC methodology, we discern two categories of models.

###### Demographic Models

Each of the four models models describes the subdivision of an ancestral population into two daughter populations (Fig. S7-A). The three populations have independently assigned effective population sizes. The differences between these four models concern the historical patterns of gene flow between two divergent populations, as depicted in figure S7. These models encompass continuous migration (CM), and secondary contact (SC), strict isolation (SI) and ancient migration (AM) :

- models with ongoing migration
  - continuous migration (CM)
  - secondary contact (SC)
- models with current isolation
  - strict isolation (SI)
  - ancient migration (AM)

Notably, the former two models entail ongoing gene flow between the two populations, while the latter two do not. Models with past (AM) or recent (CM and SC) migration assume gene flow between sister populations/species in both directions, at two independently assigned rates.

###### Models of Linked Selection

Effects of linked selection have been taken into account using a genomic model that encompasses: (a) heterogeneous effective population size across the genome (*hetero. N_e_*), which closely approximates the influence of background selection by down-scaling *N_e_* (*95*); and/or (b) heterogeneous migration rate across the genome (*hetero. M*) to account for the effects of selection against hybrids (*96*). The modeling framework employed in this study does not consider the effects of positive selection on linked loci (i.e., genetic hitchhiking).

Within the *hetero. N_e_* genomic model, the variable effective size among loci is assumed to conform to a re-scaled Beta distribution. In essence, all populations share a common Beta distribution with two shape parameters drawn from uniform distributions. However, each population is independently re-scaled by distinct *N_e_* values, which are drawn from uniform distributions. Conversely, the *homo. N_e_* genomic model assumes that all loci from the same genome share the same effective population size, and this parameter is independently estimated in all populations. This homogeneous model implies that the genomic landscape remains unaffected (or is uniformly affected) by background selection.

The *hetero. M* genomic model implements local reduction of gene flow in the genome. Variation in migration rates among loci is thus modeled by employing a bimodal distribution where a proportion of loci, drawn from a uniform distribution in ]0-1[, is linked to barriers (i.e., *N_e_.m* = 0), while the loci unaffected by species barriers are associated to an effective migration rate *N_e_.m* drawn from a uniform distribution. In the *homo. M* model, a single migration rate *N_e_.m* per direction is universally shared by all loci in the genome.

Subdivisions of the four demographic models (CM, SC, SI and AM) into various genomic submodels were made to accommodate for the effect of linked selection. Heterogeneity in effective population size was a universal consideration across all four models, while heterogeneity in migration rate was specifically accounted for in models exhibiting gene flow (i.e., CM, AM, and SC). Therefore, the SI model was divided into two submodels (*homo. N* or *hetero. N*), while the AM, CM, and SC models were divided into four submodels:

1. *homo. N_e_* and *homo. M*
2. *homo. N_e_* and *hetero. M*
3. *hetero. N_e_* and *homo. M*
4. *hetero. N_e_* and *hetero. M*

For a comprehensive description of all prior distributions employed in this study, please refer to Section A.4.5.

##### A.4.4 Summary statistics

ABC is a statistical inferential approach based on the comparison of summary statistics derived from simulated and observed datasets (*97*). We present a comprehensive description of the statistics computed within our framework. Previous studies have demonstrated the effectiveness of these statistics in statistically distinguishing demographic models with and without ongoing migration (*14, 22, 89*). The following summary statistics are calculated for each locus:

- The number of bi-allelic polymorphisms in the alignment including all sequenced copies in the 2 species/populations
- Pairwise nucleotide diversity *π* (*98*)
- Watterson’s *θ* (*99*)
- Tajima’s *D* (*100*)
- The proportion of sites displaying fixed differences between the populations/species (*S_f_*)
- The proportion of sites featuring polymorphisms exclusive to a specific population/species (*S_xA_* and *S_xB_*)
- The fraction of sites with polymorphisms shared between the two populations/species (*S_s_*)
- The number of successive shared polymorphic sites
- Raw divergence *D_xy_* between the two populations/species (*101*)
- Net divergence *D_a_* between the two populations/species (*101*)
- Relative genetic differentiation between the two populations/species quantified by *F_ST_* (*102*)

For the ABC analysis, we used the means and variances of these statistics calculated over all the available loci. Additionally, we utilize the joint Site Frequency Spectrum (jSFS (*103*)) to summarize the data, specifically capturing the count of single-nucleotide polymorphisms (SNPs) where the minor allele occurs in each bin covering the jSFS. Because of the absence of outgroup lineages, jSFS were folded. Singletons are deliberately excluded from the jSFS to mitigate potential inference biases arising from sequencing errors. Each of the non-excluded bin of the jSFS is used as a descriptive statistics in the ABC analysis.

We supplement this set of summary statistics with measures taken on all the loci:

- Pearson’s correlation coefficient for *π* between species
- Pearson’s correlation coefficient for *θ* between species
- Pearson’s correlation coefficient between *D_xy_* and *D_a_*
- Pearson’s correlation coefficient between *D_xy_* and *F_ST_*
- Pearson’s correlation coefficient between *D_a_* and *F_ST_*
- Proportion of loci with both *S_s_* and *S_f_* sites
- Proportion of loci with *S_s_* sites but no *S_f_*
- Proportion of loci without *S_s_* sites but with *S_f_*
- Proportion of loci with neither *S_s_* nor *S_f_* sites

The summary statistics obtained from both the empirical data sets (i.e., plants and animals) and the data sets simulated under the demographic models (Fig. S7) were calculated with the same scripts implemented in DILS.

##### A.4.5 Configuration file

DILS was run using the following parameter values:

- mu = 7.31 × 10^−9^
- useSFS = 1
- barrier = bimodal
- *max_N_tolerated* = 0.25
- *L_min_* = 10
- *n_min_* = 4
- *rho_over_theta* = 0.2
- uniform prior for *N_e_* between 0 and *N_e,max_* individuals
- uniform prior for *T_split_* between 0 and *T_max_* generations
- uniform prior for migration rate *N_e_.m* between 0 and 10 migrants per generations

Where:

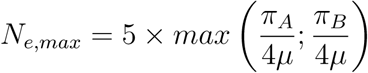

*π_A_*and *π_B_*being the Tajima’s *θ* (*98*) for species A and B respectively (for a given pair).

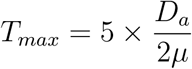

*D_a_*being the net divergence (*101*).

##### A.4.6 Returned quantities

At the end of the analysis, DILS returns the posterior probability of ongoing migration *versus* of current isolation. The probability of ongoing migration corresponds to the relative probability of all models including ongoing migration (Secondary Contact, Continuous Migration) and their sub-models (heterogeneity and genomic homogeneity for migration and effective size); while the probability of current isolation corresponds to all models and sub-models with current isolation (Strict Isolation, Ancient Migration). These quantities are used to produce the relationships between the net divergence and the posterior probability of migration (Fig. S8). For each pair of populations/species, three statuses are then assigned:

1. Strong support for genetic isolation: we identify strong statistical support for genetic isolation when our ABC framework yields a posterior probability *P*_mig_ *<* 0.1304. This threshold was empirically determined by the robustness test conducted in (*14*), where the robustness *R*_mig_ was calculated as:

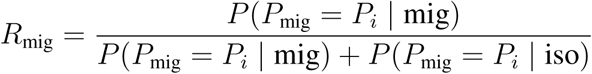 where:

- *P_i_*: the posterior probability attributed by our ABC framework to migration to a given simulated dataset.
- *P* (*P*_mig_ = *P_i_* | mig) is the probability that a dataset simulated under a model with ongoing migration is correctly inferred as a model with migration by our ABC approach, given a posterior probability for migration of *P_i_*.
- *P* (*P*_mig_ = *P_i_* | iso) is the probability that a dataset simulated under a model with current isolation is wrongly inferred as a model with migration by our ABC approach, given a posterior probability for migration of *P_i_*.
2. Strong support for ongoing migration: strong statistical support for ongoing migration is indicated when the posterior probability *P*_mig_ *>* 0.6419, also empirically determined in (*14*).
3. Ambiguity: statistical ambiguity, denoting situations where our ABC framework does not strongly support either migration or isolation, i.e, when the risk of assigning an analysed pair to a wrong status is greater than 5%.

Pairs for which support was inconclusive were excluded from further analysis. The remaining pairs were categorized either as exhibiting ‘migration’ or ‘isolation,’ as illustrated in Figure 1-A, allowing the ‘migration’ status to be treated in a logistic regression (see section A.5).

#### A.5 Logistic regression

To study speciation dynamics, we examine reduction in the proportion of plant or animal pairs receiving strong support for models with migration as a function of time (measured here by the net molecular divergence). For this purpose, we modeled **Y***_i_* (the binary status ‘isolation’ or ‘migration’ best fitting the data) as a function of **X***_i_* (the average net genomic divergence) by using a generalized linear model (GLM) *via* a linked binomial function:

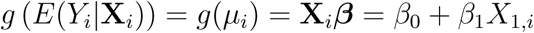

where *β*_0_ represents the intercept and *β*_1_ the coefficient reflecting the effect of genomic divergence on the isolation/migration status coded as 0 and 1, respectively. The fitted model is used to predict *p_i_*, the proportion of pairs of populations/species that are currently connected by gene flow (migration status) for a given level of divergence **X***_i_*.

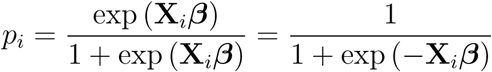

Reversely, we can determine the divergence level **X** for which a given proportion *p_i_* of pairs are connected by gene flow:

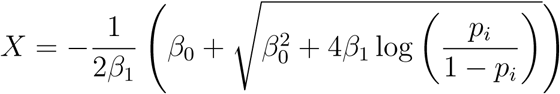

We are interested in comparing the inflection point, i.e, the level of divergence above which more than 50% of species pairs are genetically isolated, between plants and animals. Thus, for a given fitted model, this point corresponds to a divergence level 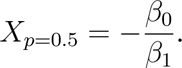

The log-likelihood function *ℓ* of the migration/isolation status **Y** given the average net molecular divergence **X** is then obtained to evaluate the fit of a model to the observed data:

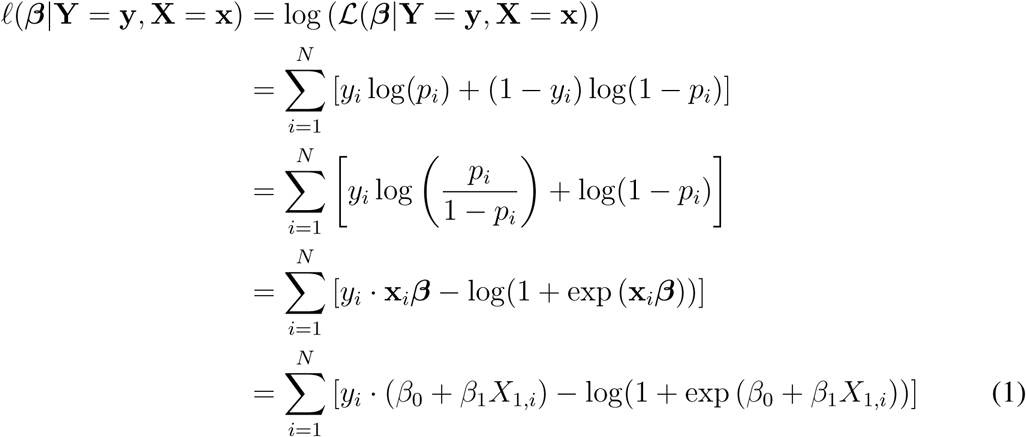

The sigmoid of plants can then be tested against that of animals to determine whether plants and animals share the same dynamics of reproductive isolation accumulation. For this purpose, three models are fitted and associated to log-likelihood *ℓ*:

1. *M*_0_: both plants and animals share the same logistic relationship between **X***_i_* and **Y***_i_*.
2. *M_plants_*: model fitted to the plants data only.
3. *M_animals_*: model fitted to the animals data only.

Thus, for *M*_0_ we fitted a GLM to the entire dataset comprising both plants and animals, after having retained only demographic inferences for which the ABC analysis produced strong statistical support for ongoing migration or current isolation, following the test of robustness applied in Roux et al. (*14*). In that sense, pairs of plants and animals with ambiguous support for isolation or migration were excluded from all GLM regressions. The log-likelihood *ℓ*(*M*_0_) was then estimated for the whole dataset comprising both plants and animals by using formula **1** where:

- ***β*_0_** and ***β*_1_** represent for *M*_0_ the coefficient of the model fitted to the whole plants and animals dataset by using the **glm** function (family = ‘binomial’) implemented in `R`.
- **X**_1_*_,i_* represents the series of observed net divergence values.
- **y***_i_* represents the series of inferred isolation/migration status.

For *M_plants_* and *M_animals_*, we fitted a GLM model only to data from the corresponding kingdom. We then estimated the log-likelihoods *ℓ*(*M_plants_*) and *ℓ*(*M_animals_*) as for *M*_0_.

Finally, we conducted a comparison between the log-likelihood *ℓ*(*M*_0_) and the combined log-likelihood *ℓ*(*M_plants_*)+*ℓ*(*M_animals_*), which is derived from the summation of log-likelihoods obtained by fitting independent models to each respective kingdom. The significance of the difference between *ℓ*(*M*_0_) and *ℓ*(*M_plants_*) + *ℓ*(*M_animals_*) was evaluated using a permutation-based approach. Specifically, the absolute difference between *ℓ*(*M*_0_) and *ℓ*(*M_plants_*) + *ℓ*(*M_animals_*) was compared to a null distribution generated from 10,000 random permutations of the data. The *P*-value corresponds to the proportion of permutations in which the absolute difference exceeded the observed value (Table S2).

#### A.6 Testing for a phylogenetic effect

To control for the variation in the number of pairs between genera, we carried out 1,000 animal-plant comparisons as for Fig. 1 but by randomly selecting a single pair per animal and plant genus (Fig. S9-A). Over these 1,000 sub-samples, the relative positions of the sigmoids were compared *via* the inflection points 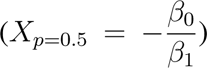 of the models fitted to the plant *versus* animal sub-samples. We consistently find that the inflection point occurs at lower levels of divergence in plants than in animals (Fig. S9-B). However, the difference in likelihood between the global model and the combined likelihoods of the plant and animal models is not significant in 7% of the permutations (Fig. S9-C; blue bars). This lack of significance is attributed to the sub-sampling process, which substantially reduces the number of data points contributing to the likelihood calculations, thereby decreasing the power to discriminate effectively between models. Nevertheless, the distance between the plant and animal inflection points was significantly greater than expected by random chance in 100% of the permutations (Fig. S9-C; orange bars).

#### A.7 Testing for a sequencing technology effect

Out of the total dataset comprising 280 pairs of plants and 61 pairs of animals, 183 plant pairs and 54 animal pairs exhibited high robustness in model comparison and passed the goodness-of-fit test. These retained datasets encompass a diversity of sequencing methodologies. Specifically, within plants, among the 183 retained pairs: 81 pairs were acquired through RAD-sequencing, 70 pairs through RNA-sequencing, and 32 pairs through whole genome sequencing. In the case of animals: 46 pairs were derived from RNA-sequencing, while 8 pairs were the result of Sanger sequencing. To assess the potential influence of sequencing techniques, we determined whether the observed differences in dynamics between plants and animals, as previously reported for the entire dataset, remained consistent when considering only the data generated exclusively through RNA sequencing. This choice was motivated by the fact that RNA-sequencing is the sole sequencing technique shared by both biological kingdoms under study. By retaining only the data from RNAseq, we maintain a statistically significant support for a more rapid cessation of gene flow in plants than in animals (*P*-value*<* 0.0001; Table S3).

#### A.8 Testing factors influencing speciation dynamics within plants

To investigate factors that may influence the dynamics of reproductive isolation in plants, we focused on two life-history traits: (i) growth form, specifically comparing herbs and trees (Section A.8.1), and (ii) mating systems by comparing different selfing rates (Section A.8.2). The first comparison is motivated by the typically greater pollen dispersal observed in trees compared to annual plants, leading to reduced nuclear genetic differentiation within tree populations (*104, 105*). For the effect of the mating system, we limited our analysis to a comparison between species in our sample with the lowest selfing rates and those with the highest. However, it is important to note that our sample is not representative of the full diversity of selfing rates found in plants (*106*), as it is biased towards high-outcrossing species (Fig. S12). The theoretical effect of selfing rate on the evolution of reproductive isolation remains unclear and appears contradictory (*107*). On one hand, higher selfing rates reduce gene flow, decrease effective recombination, and increase the strength of genetic drift, which can facilitate the fixation of deleterious mutations that may act as barriers when compensated. On the other hand, reduced efficacy of selection in high selfing species can mitigate intragenomic conflicts, potentially leading to fewer interspecific incompatibilities.

For these analyses, we focused on the plant dataset, subdivided into different groups based on the specific comparisons performed for each tested factor. We first conducted a comparison based on life form (herbs *versus* trees), followed by three comparisons to assess the effect of selfing rate (selfers *versus* outcrossers, selfers *versus* mixed systems, and outcrossers *versus* mixed systems).

Each comparison was performed similarly to the plant-*versus*-animal comparison: we first fitted a global model on the entire dataset and then tested whether the sum of the likelihoods of the models fitted on each subgroup was significantly greater than that of the global model.

##### A.8.1 Effects of plant life forms

The plant dataset was divided into four categories based on their life form:

- **Liana** (e.g., *Actinidia*, *Nepenthes*)
- **Herb** (e.g., *Arabis*, *Dactylorhiza*, *Helianthus*, *Hibiscus*, *Isoetes*, *Lupinus*, *Pitcairnia*, *Pulmonaria*, *Rhodanthemum*, *Senecio*, *Silene*, *Phlox*)
- **Tree** (e.g., *Ficus*, *Laccospadicinae*, *Phoebe*, *Picea*, *Populus*, *Quercus*, *Yucca*)
- **Shrub** (e.g., *Gossypium*, *Salix*, *Stachyurus*)

We performed two comparisons: first, between herbs and trees (Fig. S10-A), which showed no significant reduction in gene flow among herbs compared to trees (*P* = 0.261; Table S4). When herbs, lianas, and shrubs were grouped together (Fig. S10-B), we still did not observe a reduction in gene flow relative to trees (*P* = 0.1937; Table S4).

##### A.8.2 Effects of plant mating systems

We aimed to test whether selfing species exhibit a faster emergence of reproductive isolation compared to outcrossing species, as hypothesized due to reduced dispersal, decreased effective recombination, and a higher rate of accumulation of genetic incompatibilities. Alternatively, selfing species might experience a slower development of reproductive isolation compared to outcrossers, driven by reduced intragenomic conflicts that could otherwise act as barriers. Since we lacked selfing rate estimates from the literature for each species, we inferred these rates from the genomic data used for demographic analyses. The developed method estimates the individual inbreeding coefficient *F* for an individual *i*, providing the selfing rate *s* under the assumption of an equilibrium population. This is achieved using a maximum likelihood approach based on the observed genotypes and allele frequencies at polymorphic loci.

###### Definitions and Notations

- *s* is the equilibrium selfing rate, where *s* ∈ [0, 1].
- *p* is the allele frequency of the alternative allele at a given site.
- *q* = 1 − *p* is the frequency of the reference allele.
- *F* is the inbreeding coefficient, which is related to the selfing rate *s* at equilibrium by:

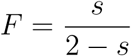

###### Genotype Probabilities

For a given site, the probabilities of observing the genotypes are modeled using *p*, *q*, and *F* :

- Probability of observing genotype “11” (homozygous for the alternative allele):

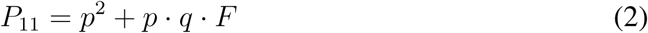
- Probability of observing heterozygous genotype “10” or “01”:

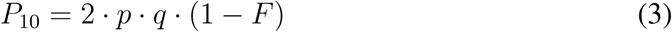
- Probability of observing genotype “00” (homozygous for the reference allele):

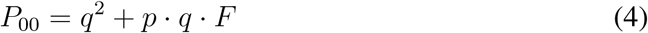

###### Log-Likelihood Calculation

For an individual *i*, the log-likelihood of the equilibrium selfing rate *s* explaining *F* is computed by summing the log-probabilities of the observed genotypes across all polymorphic sites:

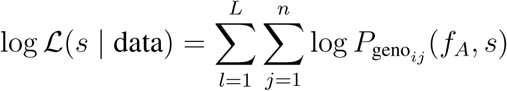

Where:

- *L* is the number of polymorphic loci (i.e, loci containing at least one polymorphic site).
- *n* is the number of polymorphic sites within each locus.
- *P*_geno_ is the probability of the observed genotype at site *j* for individual *i*, calculated based on equations (2-4).

###### Selfing Rate Estimation

We evaluate the log-likelihood for a range of selfing rates *s* (from 0 to 1 in steps of 0.05). The estimated selfing rate 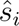 for individual *i* is the value of *s* that maximizes the log-likelihood:

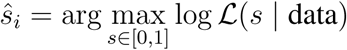

###### Numerical Stability

To avoid issues with numerical underflow when computing log-probabilities, a small constant *ɛ* = 10^−10^ is added:

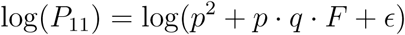

This ensures that the probabilities are always strictly positive, preventing errors in the log calculations.

###### Implementation

The entire pipeline for estimating selfing rates using the maximum likelihood approach described above has been implemented in Python. This method, along with detailed documentation and example datasets, is freely available as an open-source project on GitHub: https://github.com/popgenomics/selfing_ML.

###### Accuracy of the estimate

We evaluated the performance of our method across a range of 11 selfing rates (*s*) from 0 to 1. Genotype data were simulated using the R package `hierfstat` (*108*), which generates diploid genotypes for populations with a specified inbreeding coefficient (*F*), given a defined number of loci and individuals. The simulated datasets were then analyzed with our custom tool to estimate the selfing rate (*s*), allowing us to assess both the accuracy and the limitations of the method. We investigated the impact of varying sample sizes (2, 3, 4, 5, and 10 diploid individuals) and explored datasets consisting of 1,000 and 5,000 SNPs. Each combination of selfing rate, SNP count, and sample size was replicated 20 times.

Overall, selfing rate estimates (*s*) are substantially underestimated when the sample consists of only 2 diploid individuals (Fig. S11). Thus, for simulations with a sample size of 2 diploids and low true selfing rates (below 0.3), the inferred selfing rate is frequently estimated as zero. This bias diminishes rapidly with an increase in sample size. However, even with small sample sizes, our method reliably detects high selfing rates when the true value is elevated, although the estimates tend to be slightly underestimated. This underestimation in small samples can be attributed to the overestimation of true allele frequencies for rare alleles, which are the most common type of alleles in a site frequency spectrum. In a sample of only 2 diploid individuals, a rare allele would have a minimum frequency of 25%, which does not accurately reflect its true population frequency. This inflated allele frequency in small samples biases the selfing rate estimate downward by making the absence of homozygotes for such alleles unlikely under a non-zero selfing rate. This effect diminishes both with a moderate increase in sample size (beyond 2 individuals) and at higher true selfing rates.

###### Application to empirical plant dataset

We applied the developed method to the 135 plant species included in our study. The results revealed that the variance in selfing rates across our dataset is smaller than reported in previous dedicated studies (*106*), primarily due to the absence of strongly selfing species in our dataset (Fig. S12).

#### A.9 Geography

Geographical (geodesic) distance in meters was measured using GPS coordinates provided in the metadata when available, using the **distGeo** function in the `R` package *geosphere*. For a given pair of populations/species A and B, this distance corresponds to the distance between the two geographically closest individuals. In the case of sampled sympatric pairs, and if a single coordinate was provided by the authors for all individuals A and B, we consider a distance of 10m in line with current sampling practices to reduce relatedness. Among the 25 plant genera under examination, our review of the literature has not yielded information pertaining to the geographical origins of specimens from *Gossypium*.

To test whether the observed pattern of reduced gene flow in plants compared to animals could be merely explained by greater geographic distances between plant pairs, we performed logistic regression analyses separately for plants and animals. Using the GPS coordinates of sampled individuals, we calculated the minimum geographic distance (`distances meters`) between taxa within each pair.

We then modeled the probability of ongoing migration (`P ongoing migration`) as a function of geographic distance and net divergence (`netdiv avg`) using binomial logistic regression with a logit link function. The logistic regression results showed that geographic distance (`distances meters`) was not a significant predictor of ongoing migration for either animals (*P* = 0.688) or plants (*P* = 0.937), confirming that our conclusions are not driven by geographic factors.

#### A.10 Data availability

All assembled datasets, the reference list used for mapping, the results of demographic inference, and the R scripts for statistical analyses and figure generation are available on Zenodo under the DOI https://doi.org/10.5281/zenodo.14365924 (*51*).

### B Supplementary text

Here, we endeavor to highlight the leads that merit further exploration to unravel why plants seem to exhibit a more rapid cessation of gene flow than animals (Fig. S14).

#### Is chloroplast an additional source of barriers (Fig. S14-1)?

A hypothesis that naturally arises centers on a feature absent in animals but universally present in (non-parasitic) plants: the chloroplast. This plant-specific organelle has been shown to drive species-specific co-evolution with the nuclear genome, with the latter accumulating compensatory mutations to maintain coadaptations essential for nucleo-cytoplasmic interactions (*32*). In hybrid individuals, the presence of compensatory mutations without the co-adapted chloroplast genome, or the chloroplast genome without its corresponding nuclear compensatory mutations, can lead to developmental disruptions that significantly reduce gene flow between closely related plant species (*33*). Additionally, the generally lower mitochondrial mutation rates in plants (*34*) may accelerate speciation by allowing for compensatory selection, thereby accelerating the evolution of mitonuclear incompatibilities that contribute to reproductive isolation (*35*). Finally, the mutation rate of chloroplast is relatively low compared to the mitochondrial mutation rate observed in animals. This suggests that the theory outlined above for mitochondria could also apply to incompatibilities between the nucleus and chloroplast. Such incompatibilities would involve additional loci, further increasing the number of cyto-nuclear incompatibilities.

#### What is the role of the more prevalent selfing in plants (Fig. S14-2)?

An important characteristic of plants lies in their remarkable diversity in mating systems, particularly evident among hermaphrodites, where a broad spectrum of self-fertilization rates exists, ranging from high-selfers to high-outcrossers (*36*). Phylogenetic evidence demonstrates that the former exhibit a greater speciation rate compared to the latter (*37*). At a microevolutionary level, interspecific gene flow is higher within pairs of outcrossers compared to pairs comprising selfers (*39*). There are several reasons that may explain these patterns in selfers: reduced number of available ovules for cross-fertilization, reduced resource allocation to pollen production (*109*), autogamous F1s produce more F2 than backcross genotypes suffering from greater hybrid breakdown (*110*), and their reduced effective population size and increased linkage disequilibrium facilitate the establishment of barriers between species (*38*). Thus, frequent evolutionary periods with intermediate to high selfing in plants could contribute to our observation that genetic exchange is less frequent on average between plants than animals.

#### Would dependence on an external pollinator strengthen isolation (Fig. S14-3)?

Despite the historical emphasis on pre-zygotic isolation through behavioral mate choice as a determinant of reduced interspecies gene flow in animals, it is worth noting the occurrence of specific pre-zygotic isolation mechanisms in plants too. First, phenological shifts between related plant taxa can cause allochronic isolation (*111*). Second, plant species with animal pollination can exhibit pollinator shifts between closely-related species, playing in such situations a significant role in reproductive isolation (*40*). While these shifts are unlikely to emerge in sympatry, sister plant lineages that experienced initial divergence in allopatry can undergo divergence in their pollinator interaction networks, thus contributing to a reduced hybridization rate. This effect on reproductive isolation is reflected at the macroevolutionary scale with higher diversification rates in plants which are associated with fewer pollinator species (*41*). Such dependency on specific pollinators for reproduction thus represents a layer of co-adaptation particularly present in plants, which, as with all co-adaptations and co-evolutions, forms additional targets for forming species barriers.

#### Different modalities of dispersal (Fig. S14-4)?

Plants and animals obviously differ in their modes of dispersal, with two types of propagules, pollen and seeds, passively dispersed through abiotic and biotic vectors in plants, as compared to individual (or mother-mediated dispersal for some mammals (*112*)) mobility in animals. The overall effect of such differences is that the extent of gene flow among populations within plant species is lower (and genetic differentiation is higher) on average than in animals (*28, 29*). This may result from overall differences in dispersal kernels between plants and animals, but also from the stronger stochasticity of dispersal in plants (*113*) as compared to animals (excluding passive dispersers, e.g., marine animals with a planktonic larval dispersal), which results in higher effective migration in the latter. Stronger genetic differentiation among conspecific populations may trigger the evolution of partial reproductive isolation, a phenomenon known as outbreeding depression, which is thought to be more common among plant than animal populations (*42*).

#### Is haploid selection against barriers stronger in plant hybrids (Fig. S14-5)?

Haldane’s rule stems from the empirical observation that heterogametic hybrids (XY males and ZW females, i.e., the sex carrying a single functional copy of the sex chromosome) suffer more from species barriers than homogametic ones (*114*). From this rule, it is deduced that a significant proportion of interspecies barriers are recessive (*44*). Despite the scarcity of comprehensive data on gene expression levels during haploid stages in plants and animals, available evidence suggests that animals exhibit minimal haploid selection, while a notable portion of the plant genome can be expressed in the pollen grain and pollen tube (*43*). The higher prevalence of recessive barriers being manifested in pollen rather than in sperm may affect more profoundly the fertility of plant hybrids compared to animal hybrids.

#### What are the consequences of open exposure to environmental pollen in plants *versus* internal fertilization? (Fig. S14-6)?

Fertilisation in plants is an interactive process between the pollen grain and the maternal tissues, involving several stages: adhesion and hydration of pollen grains on the stigma, germination of the pollen tube, navigation through maternal tissues and penetration into the embryo sac. For example, in *Arabidopsis*, the male gametophyte interacts with seven different tissues before contributing to fertilization (*115*). As the stigmatic papilla can be exposed to environmental pollen from many different species, each of these male-female interactions during fertilisation represents a target for selection on the female side to avoid allocating resources towards producing a seed of lower fitness. This selection pressure may reinforce the plants’ stringent response to avoid fertilisation at lower levels of divergence as compared to animals (*116*).

#### How does the expression of parental conflicts during seed development increase reproductive isolation in plants (Fig.S14-7)?

Plants often receive pollen from a varying number of different donors (*117*). This ubiquitous polyandrous reproduction gives rise to intragenomic conflicts during seed development: paternal genes strive to allocate resources preferentially to the seeds of their own offspring, potentially at the expense of unrelated seeds. Conversely, maternal genes favor allocating resources among a larger number of seeds produced as well as to its own survival (in perennial plants). This intragenomic conflict leads to coevolution between selfish genetic elements (that seek to enhance their own transmission by diverting maternal resources) and mutations (that restore a fair allocation of resources to all seeds). From these conflicts emerge species barriers (*46*) when the collection of selfish elements is placed in a hybrid genome without the restorers with which it has coevolved (and *vice versa*), causing severe negative effects on development and germination of hybrid seeds (*47*).

#### What are the role of differences in the genetic architecture of reproductive isolation?

Quantitative genetic studies investigating the genomic architecture of RI have revealed notable differences between plants and animals. In plants, reproductive isolation often involves less complex genetic interactions compared to animals, with fewer loci exhibiting very strong individual effects (*23–25, 48*). Furthermore, alleles associated with reproductive isolation in plants frequently exhibit intraspecific polymorphism, indicating that stochastic processes often maintain these alleles in populations until their potential fixation. In contrast, speciation genes in animals are more likely to show signatures of positive selection, suggesting stricter conditions for the establishment of barriers in animals than in plants (*49*).

### C Supplementary Figures

**Figure S1:**
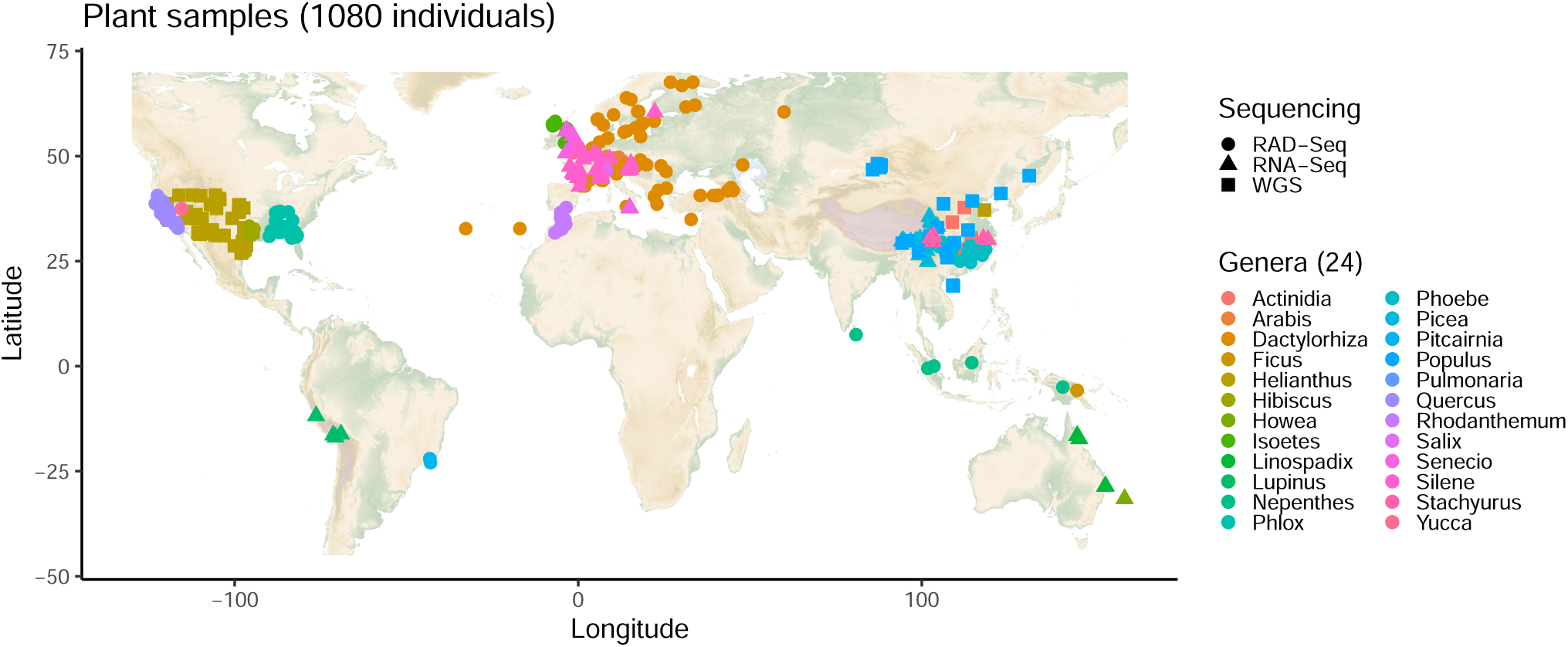
Geographical location of plant samples and sequencing methods. Each symbol represents a sequenced plant individual. The shapes correspond to the sequencing technologies used, while the colors indicate the genera.

**Figure S2:**
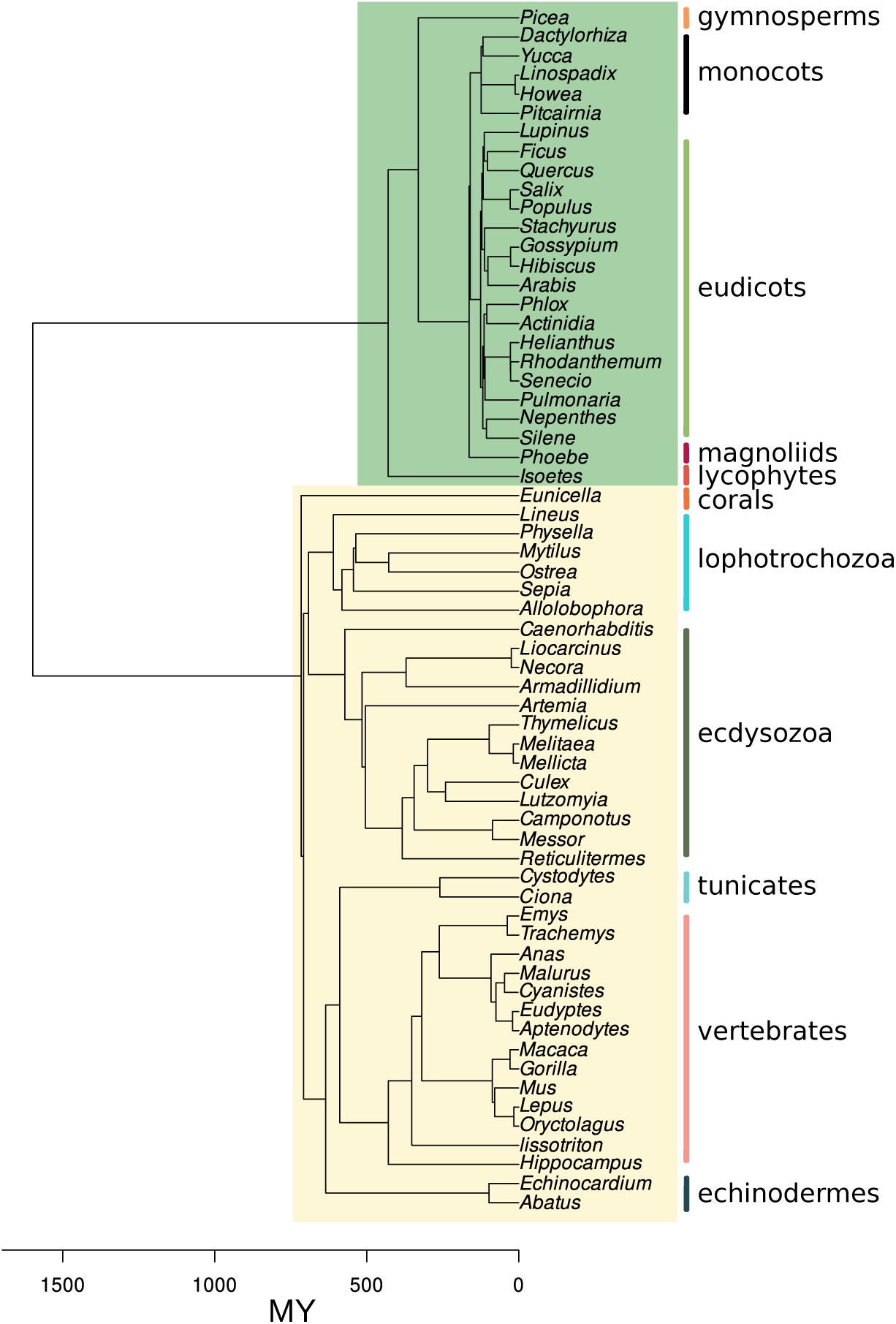
Phylogenetic relationships between species included in the current study. Plants and animals are indicated by green and yellow rectangles respectively. The scale represent the time from present expressed in million years (MY) according to `TimeTree` (*118*). Animals (yellow square) are from (*14*). Plants (green square) are included in the current study.

**Figure S3:**
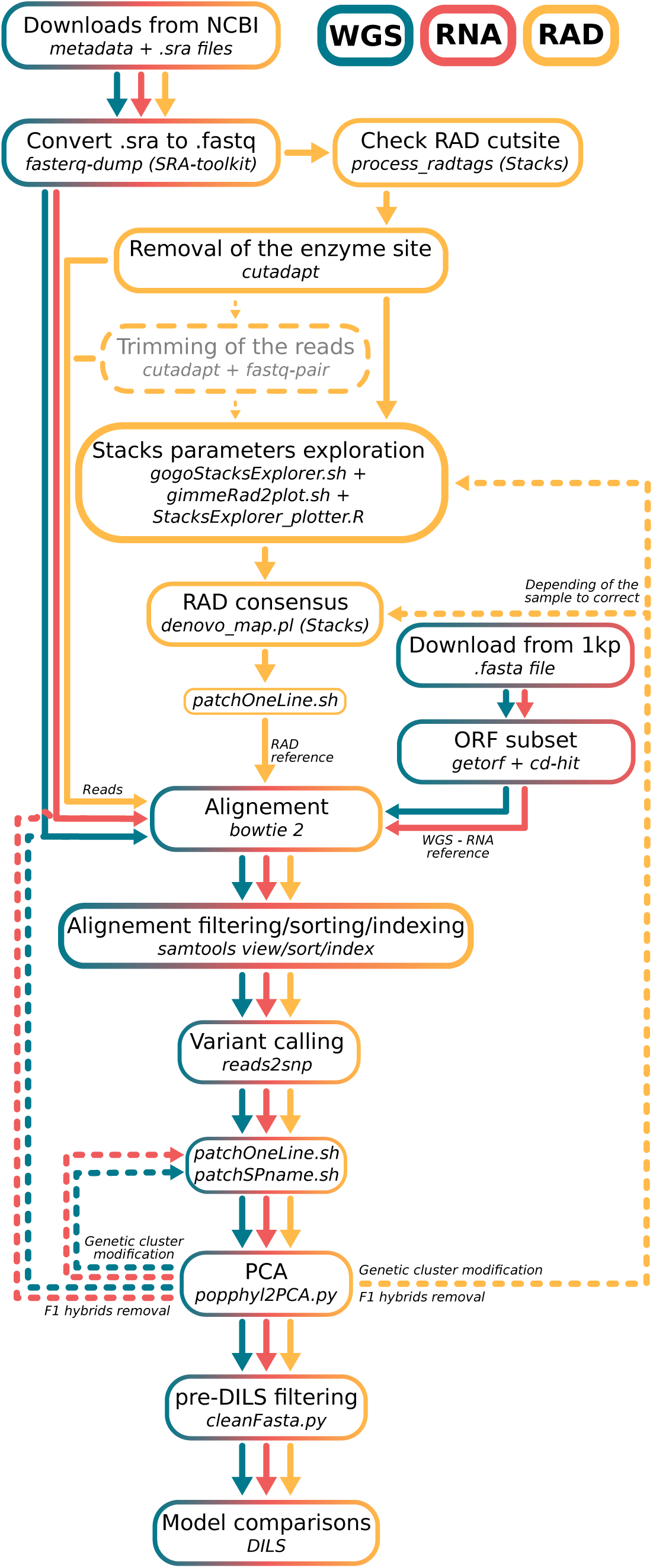
Bioinformatics steps from the raw reads to demographic analysis. Within each box, the upper line delineates an information technology procedure employed for data processing, while the lower line specifies the program or script utilized for its execution. The coloration denotes the specific sequencing technology concerned by each step.

**Figure S4:**
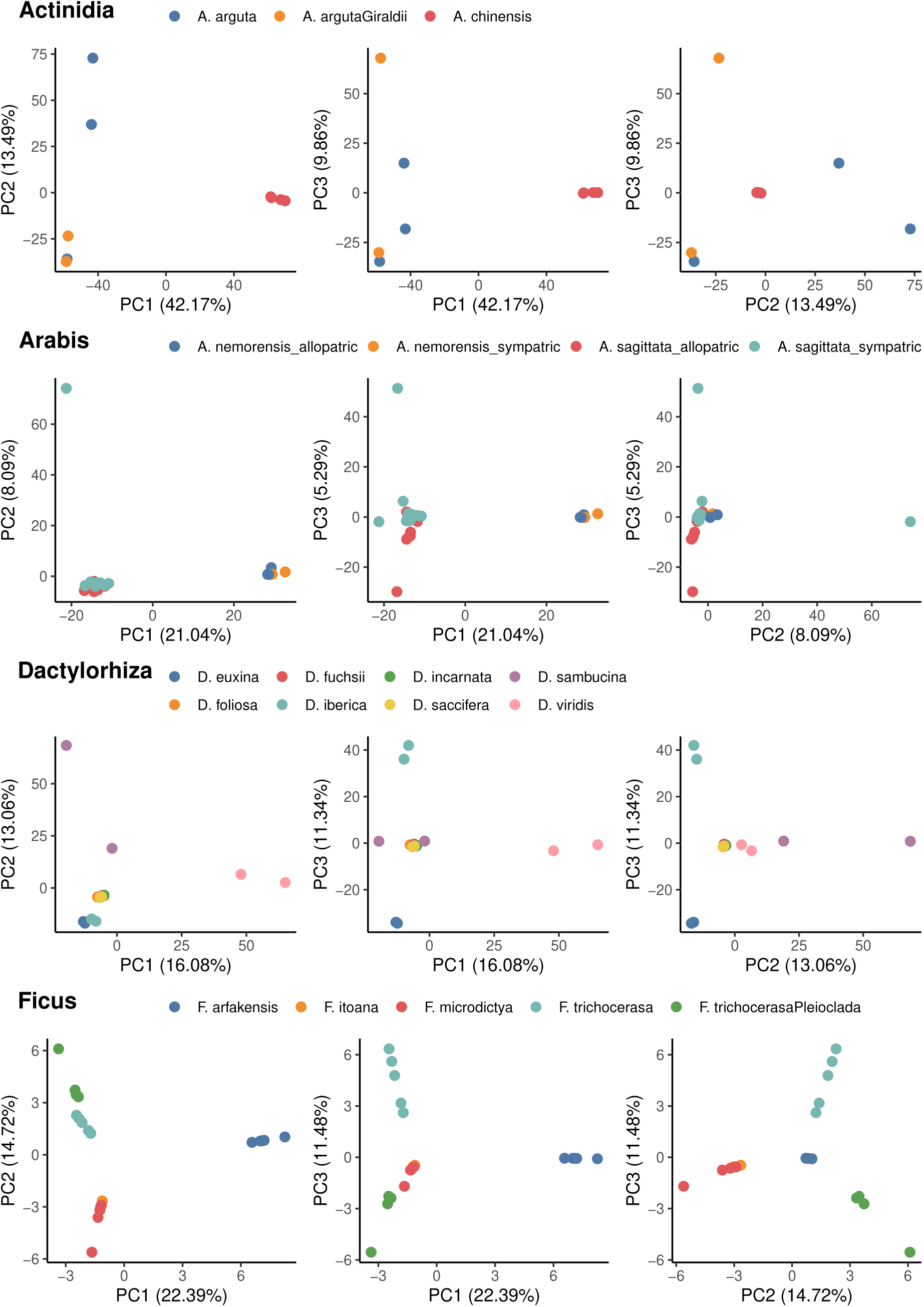
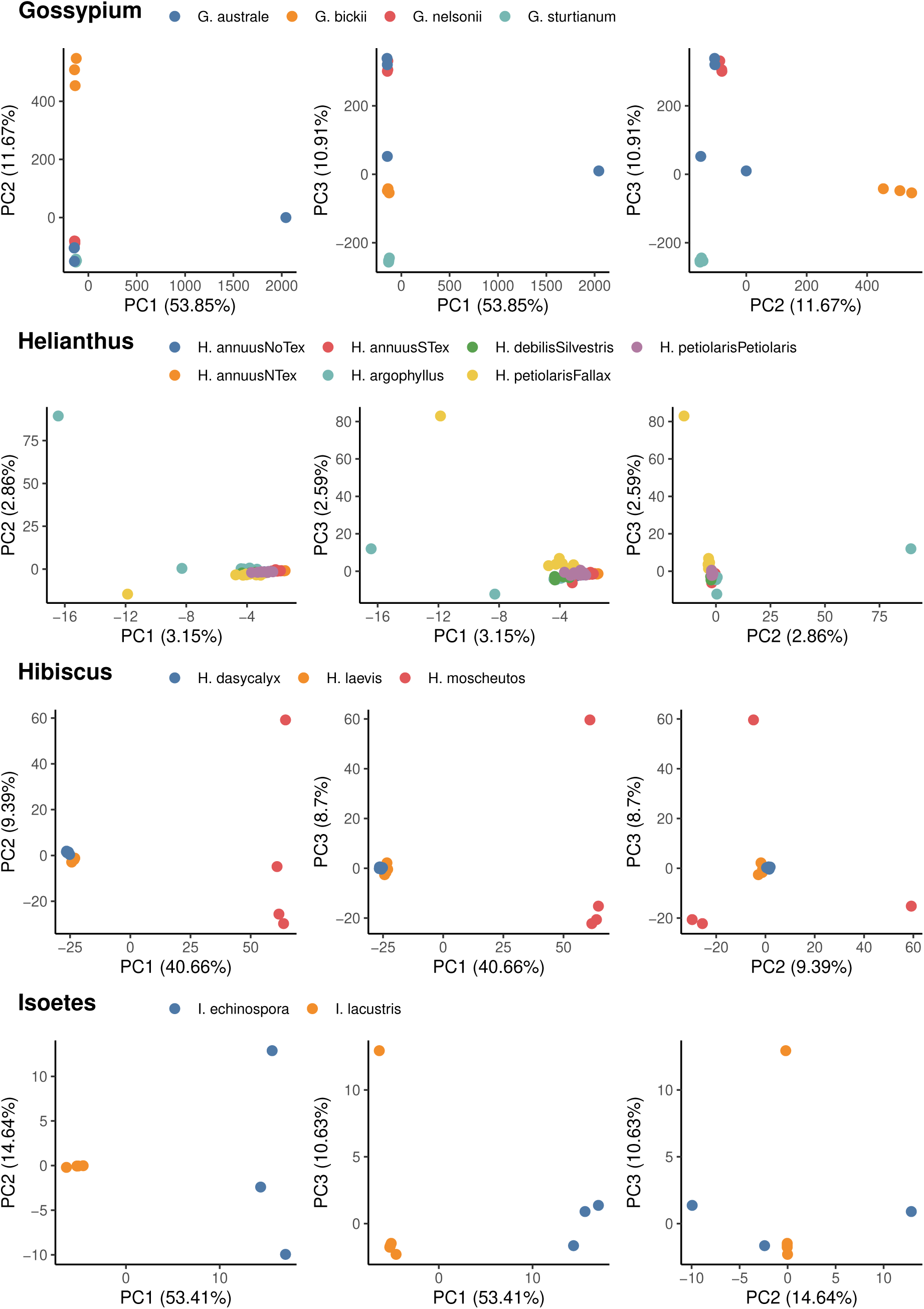
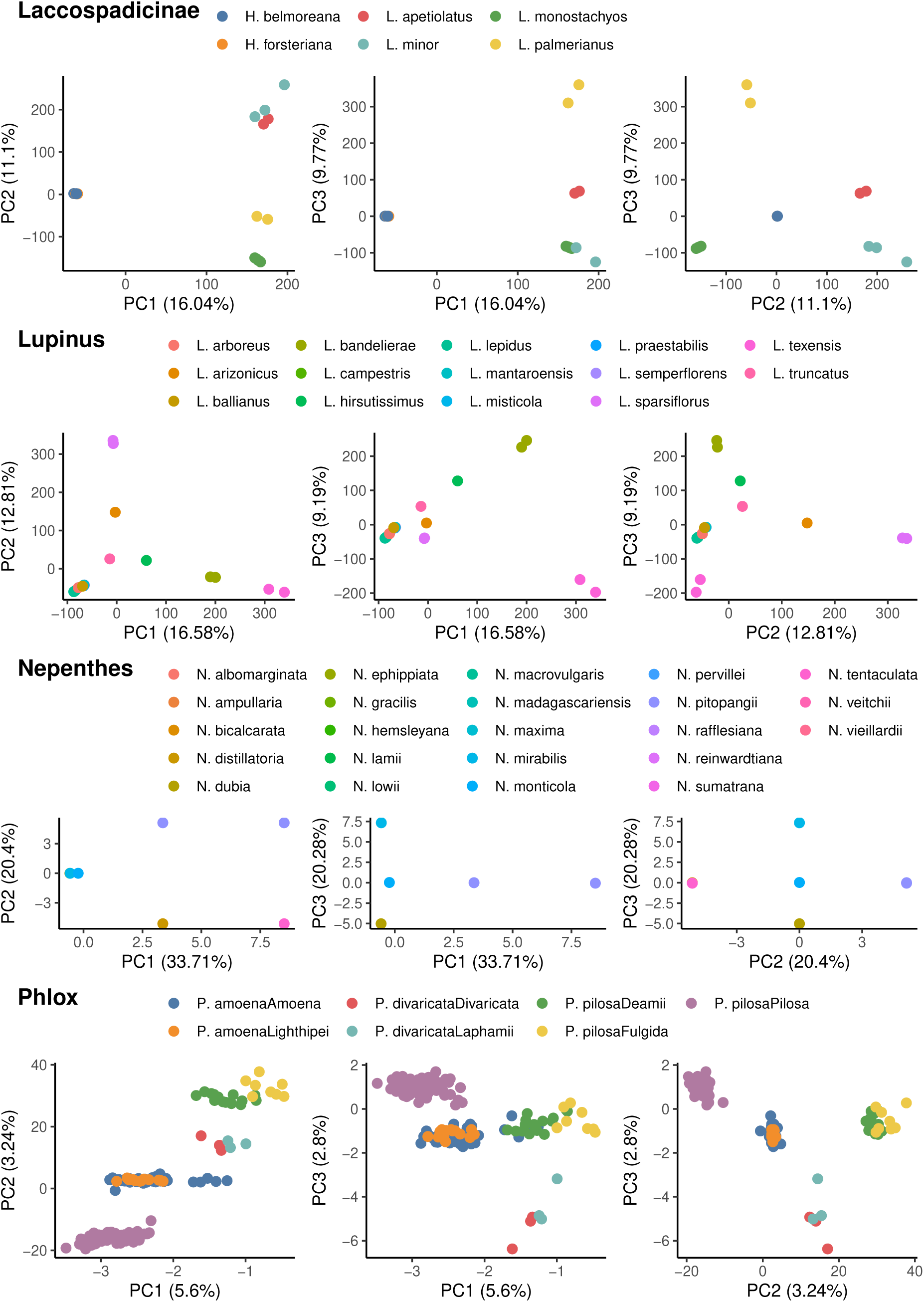
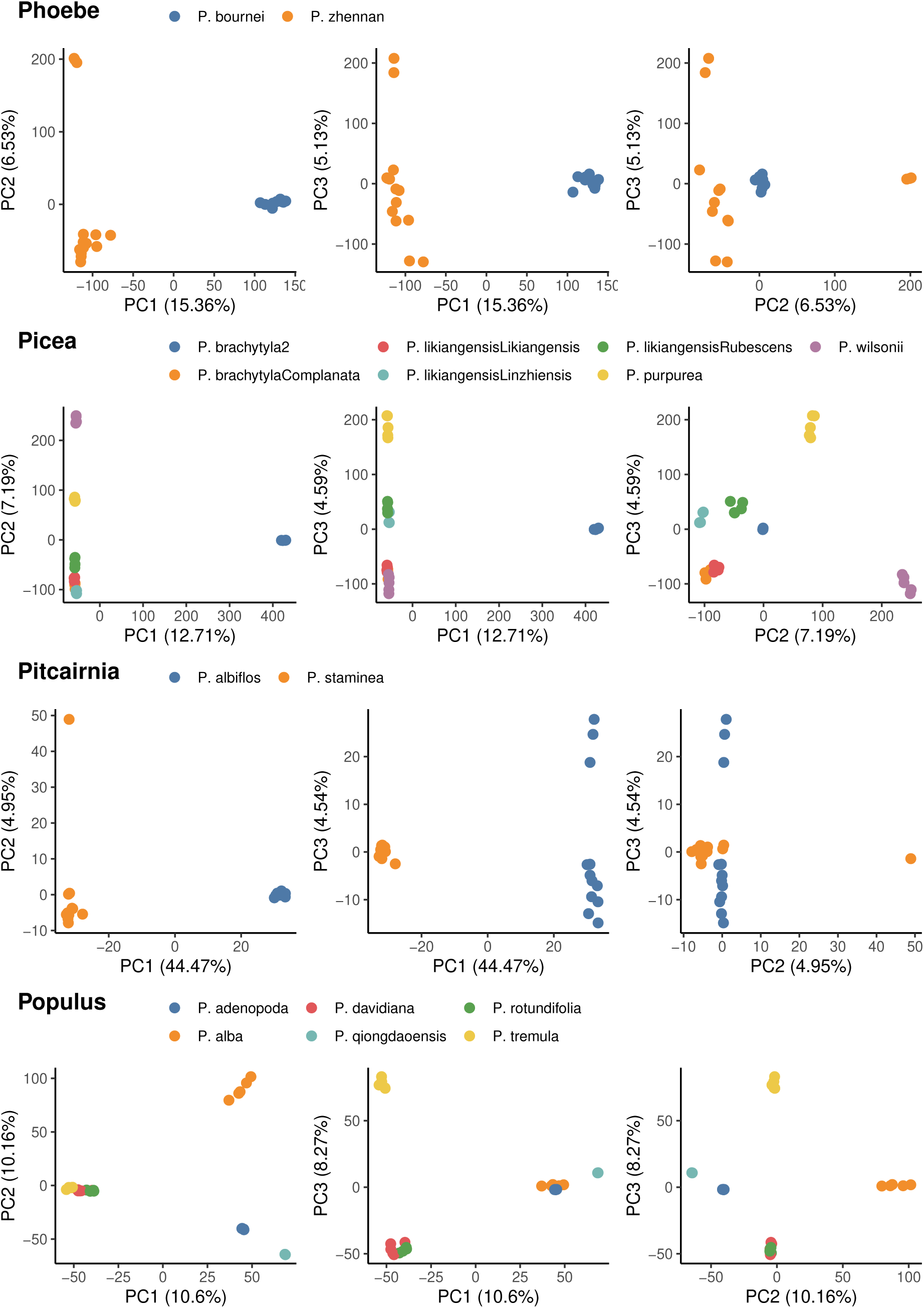
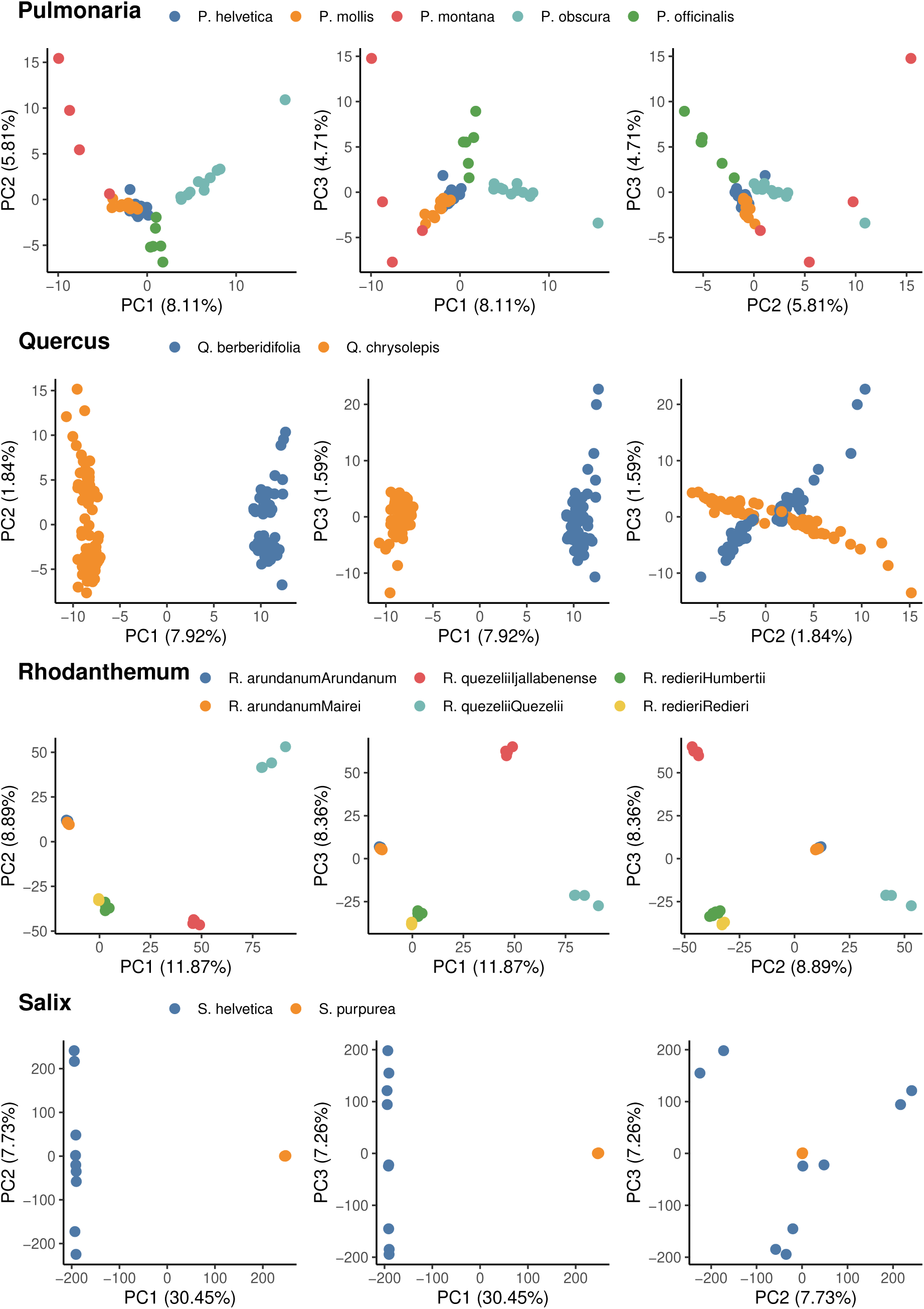
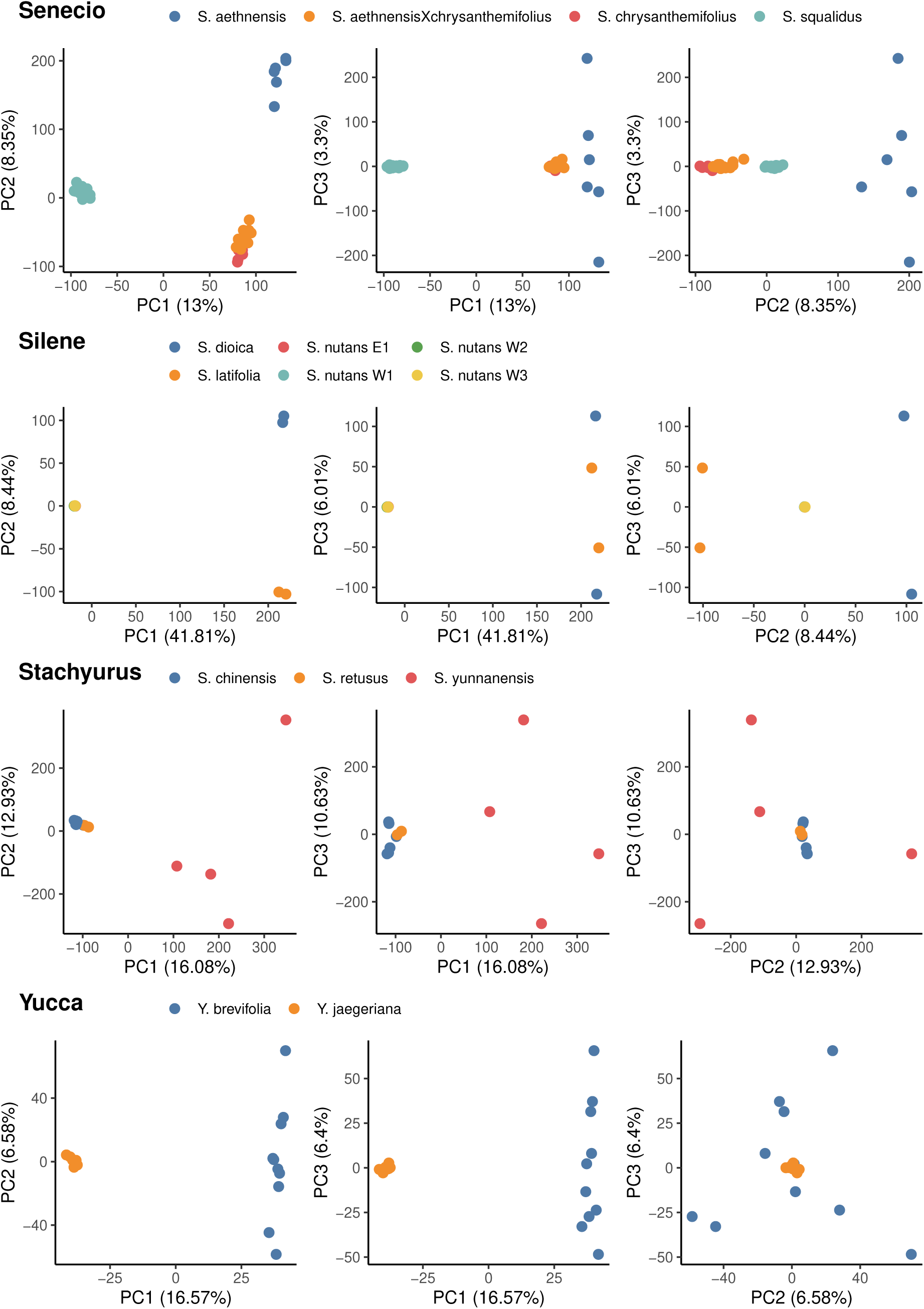
Principal component analyses on genotypes for all SNPs. Each point represents an individual. The colours represent the different populations/species named by the authors of the studies from which the data originated.

**Figure S5:**
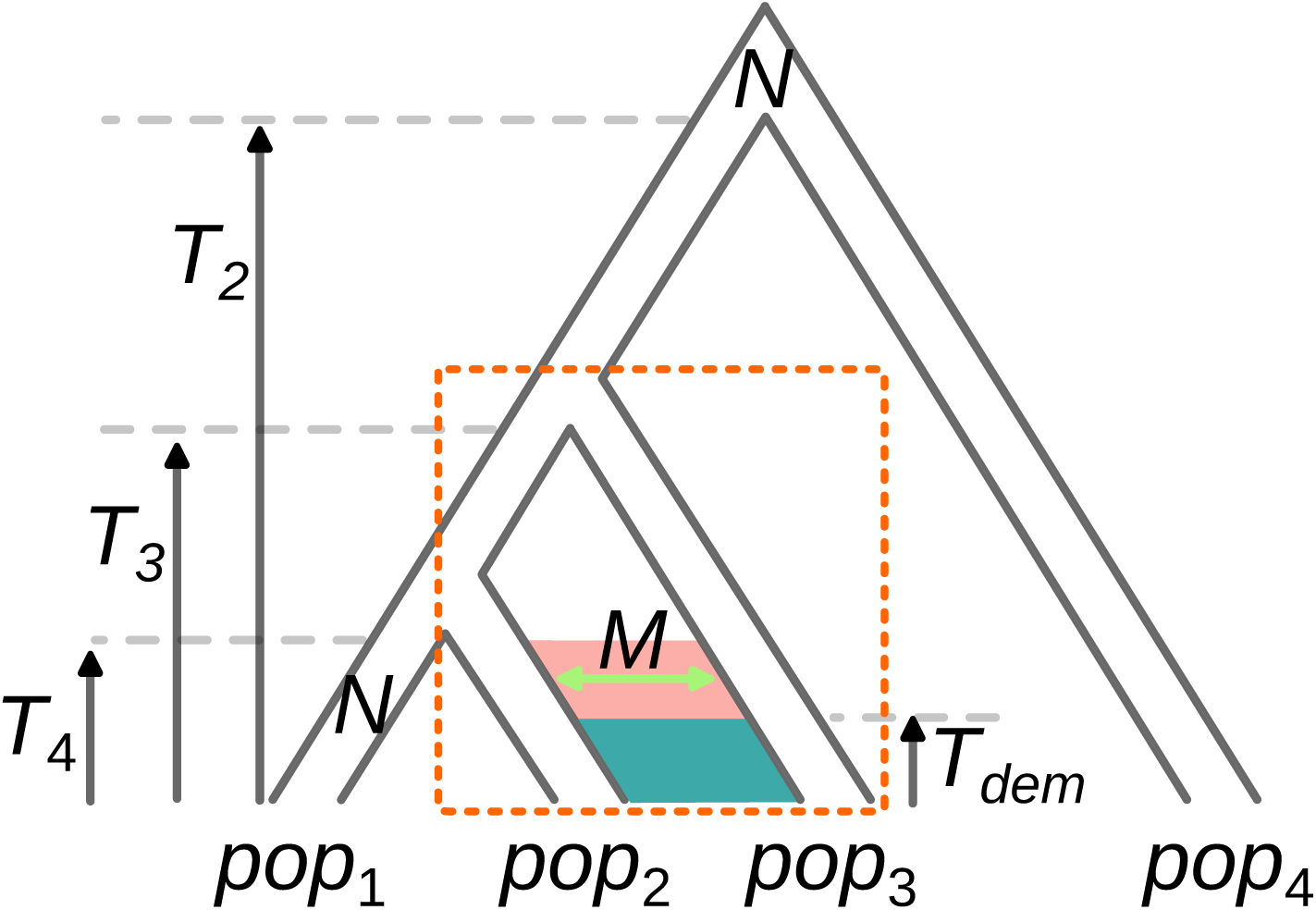
Simulated demographic model for testing introgression between populations pop_2_ and pop_3_ (dotted rectangle) using DILS and QuIBL. Parameters: *T*_4_ (speciation time between 1 and 2), *T*_3_ (speciation time between (1, 2) and 3), *T*_2_ (speciation time between ((1, 2), 3) and 4), *T*_dem_ (migration onset/end), *M* (number of migrants per generation), *N_e_* (effective population size). Scenarios: SI_4pop_ (*M* = 0 during red and blue periods), AM_4pop_ (*M >* 0 during red, *M* = 0 during blue), IM_4pop_ (*M >* 0 during red and blue), SC_4pop_ (*M >* 0 during blue, *M* = 0 during red).

**Figure S6:**
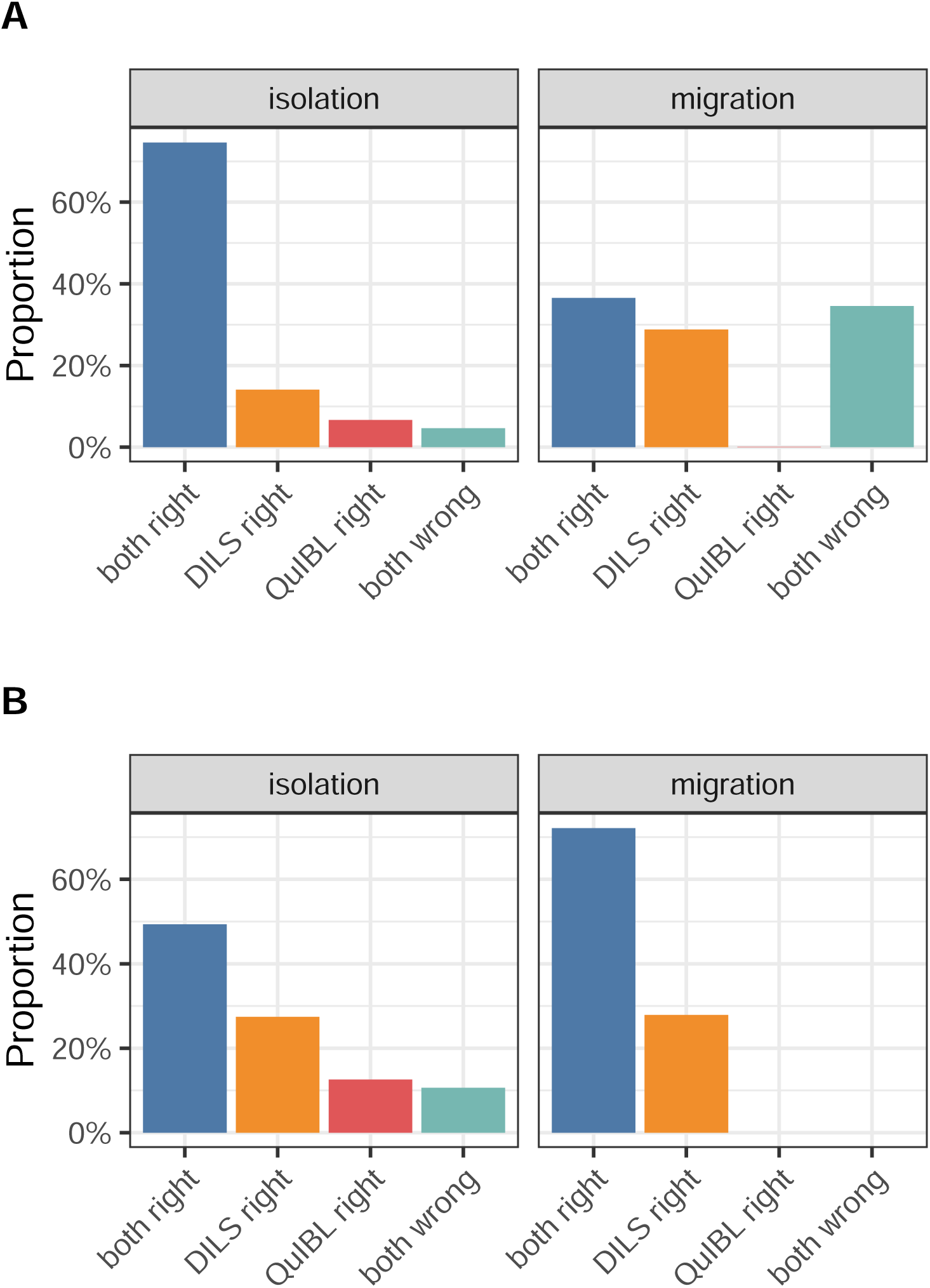
Comparison of DILS and QuIBL results on simulated datasets with migration (IM_4pop_, SC_4pop_; Fig. S5) and without migration (SI_4pop_, AM_4pop_; Fig. S5)). Colors represent the proportions of simulations where: both methods were correct (blue), only DILS was correct (orange), only QuIBL was correct (red), and both methods were incorrect (green). **A.** Results across all explored parameters. **B.** Results for parameters where more than 10% of loci are affected by migration and *N_e_.m >* 0.25.

**Figure S7:**
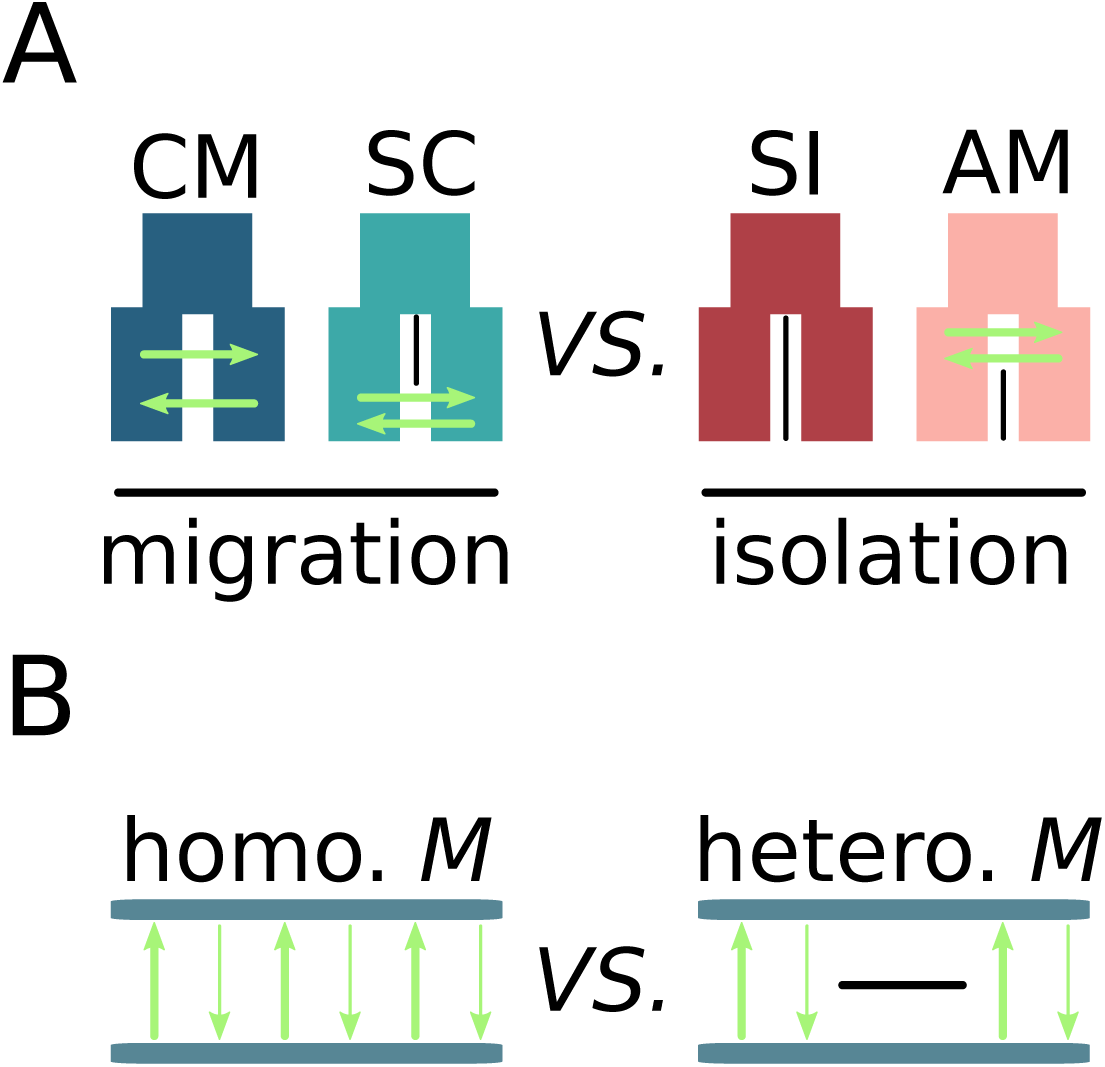
Compared models using approximate Bayesian computation (ABC). **A.** Models with ongoing migration correspond to all CM (Continuous Migration) and SC (Secondary Contact) models. Models with current isolation correspond to all SI (Strict Isolation) and AM (Ancestral Migration) models. The first step in our ABC classification is to compare the set of CM+SC *versus* SI+AM models in order to assign a migration or isolation status to each of the 341 pairs of lineages (61 animals, 280 plants) according to the computed posterior probability. **B.** Pairs of plants or animals, for which our ABC framework has provided strong statistical evidence of ongoing migration, are subsequently subjected to analysis aimed at discerning the uniformity of gene flow across the genome, whether it exhibits homogeneity (characterized by the absence of local genomic barriers) or heterogeneity (signifying genetic linkage to species barriers). The comparison between homo. *M versus* hetero. *M* was carried out using the same ABC framework as in the previous step.

**Figure S8:**
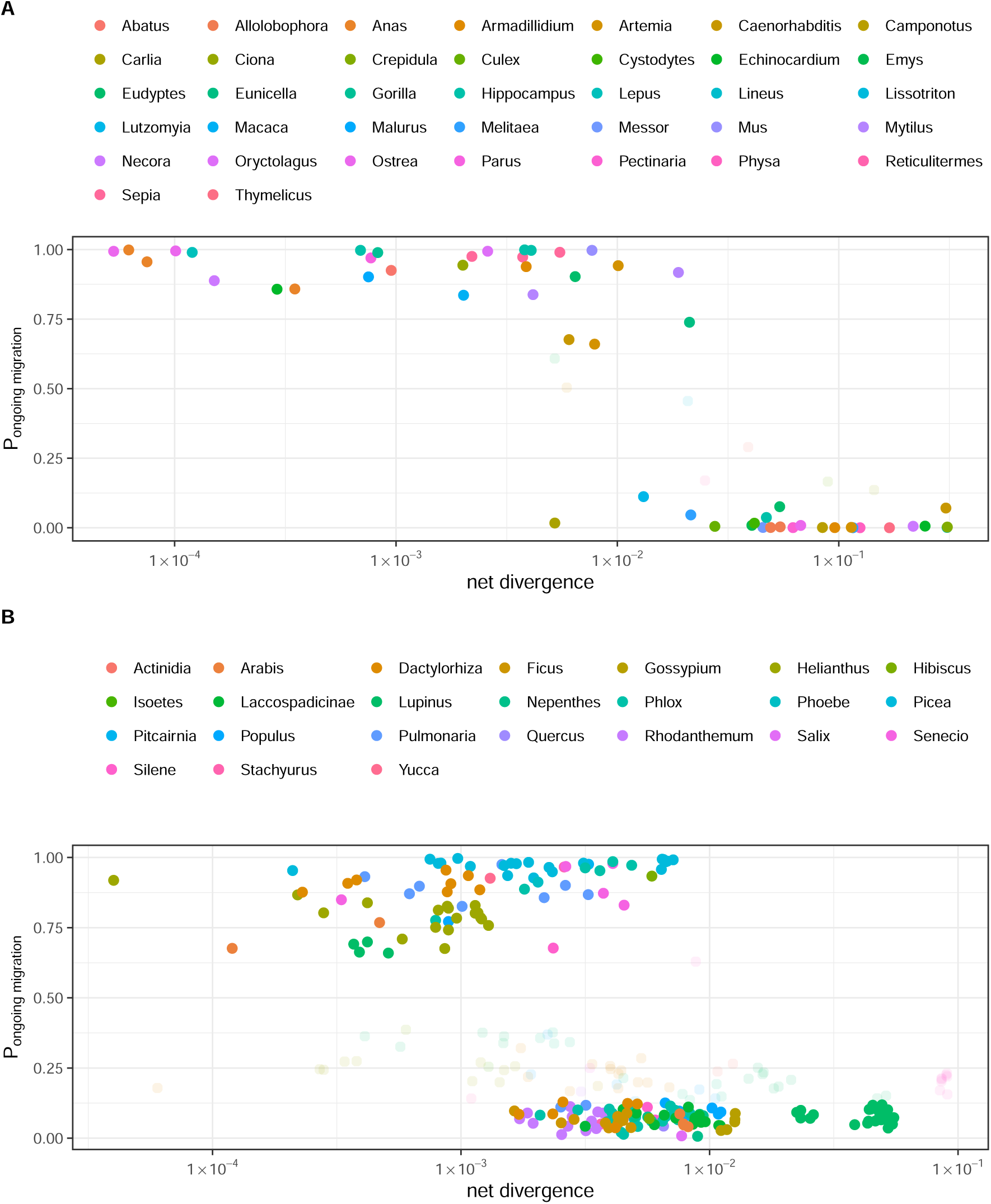
Relationship between mean net divergence and posterior probability for ongoing migration. Each point corresponds to a pair of animals (A) or plants (B). x-axis: average net divergence. y-axis: posterior probability for ongoing migration attributed by our ABC framework. Colours correspond to surveyed genera. Solid points represent pairs for which there is strong statistical evidence either supporting or rejecting the ongoing migration model, as determined by the robustness test outlined in (*14*). In contrast, transparent points indicate pairs for which the comparison between the migration and isolation models yields an inconclusive result. Pairs for which support was inconclusive were excluded from further analysis. The remaining pairs were categorized either as exhibiting ‘migration’ or ‘isolation’, as illustrated in Figure 1-A (see section A.4.6).

**Figure S9:**
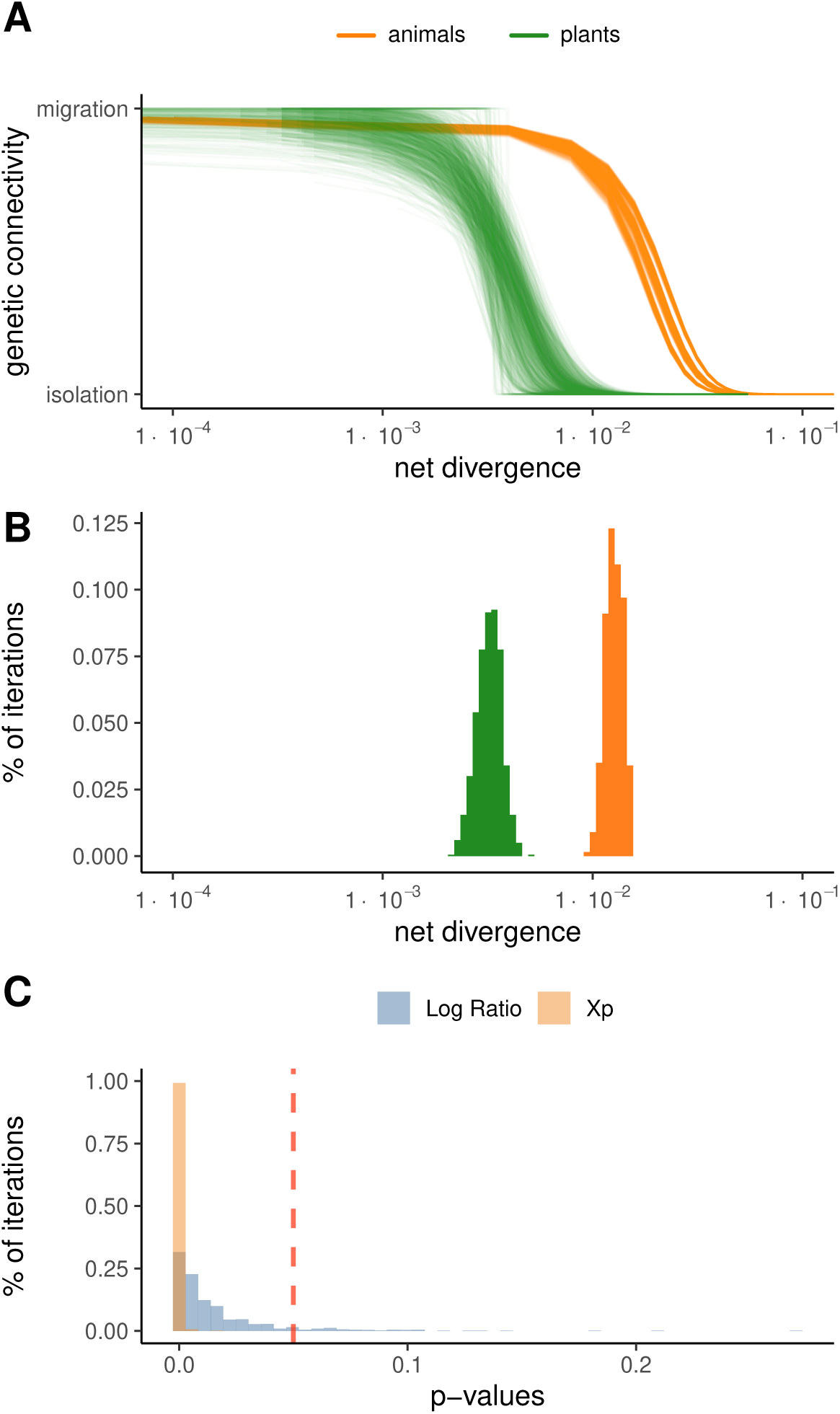
Test of the robustness of differences between plants and animals to genus effects through random subsampling. **A:** Relationship between net divergence and migration/isolation status based on random subsampling of one population/species pair per genus (plants: green; animals: orange), repeated 1,000 times. Each line represents one sub-sampling. **B:** Distribution of inflection points (*X_p_*_=0.5_) for plants and animals across 1,000 sub-samplings. **C:** *P*-value distributions from 1,000 sub-samplings: blue bars represent the log-likelihood ratio tests, and orange bars represent the tests on difference between inflection points (*X_p_*_=0.5;animals_ − *X_p_*_=0.5;plants_). The red dashed line indicates the 0.05 significance threshold.

**Figure S10:**
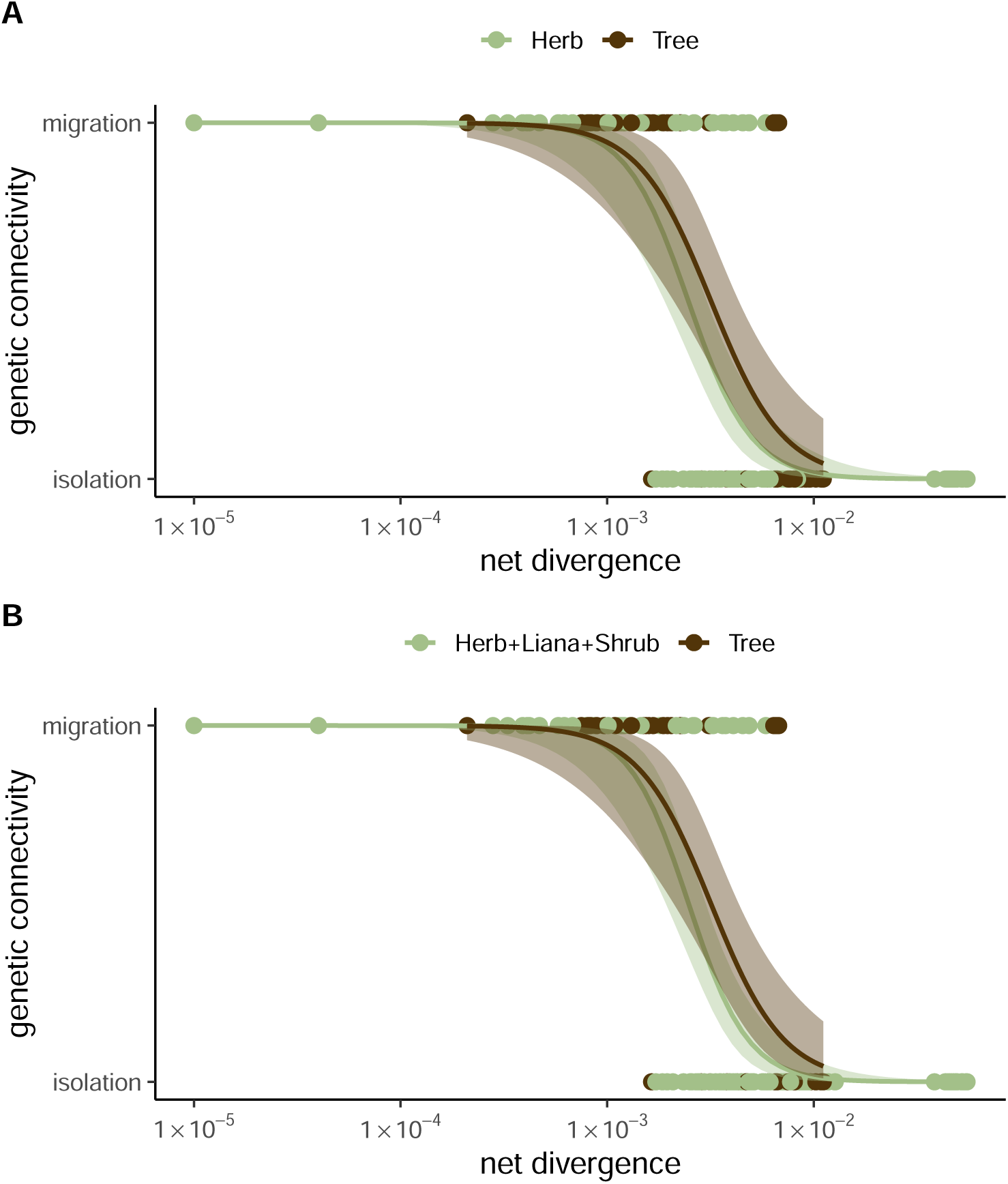
Relationship between mean net divergence and migration/isolation status across plant life forms. **A:** Comparison between herbs and trees. **B:** Comparison between herbs, lianas, and shrubs combined *versus* trees.

**Figure S11:**
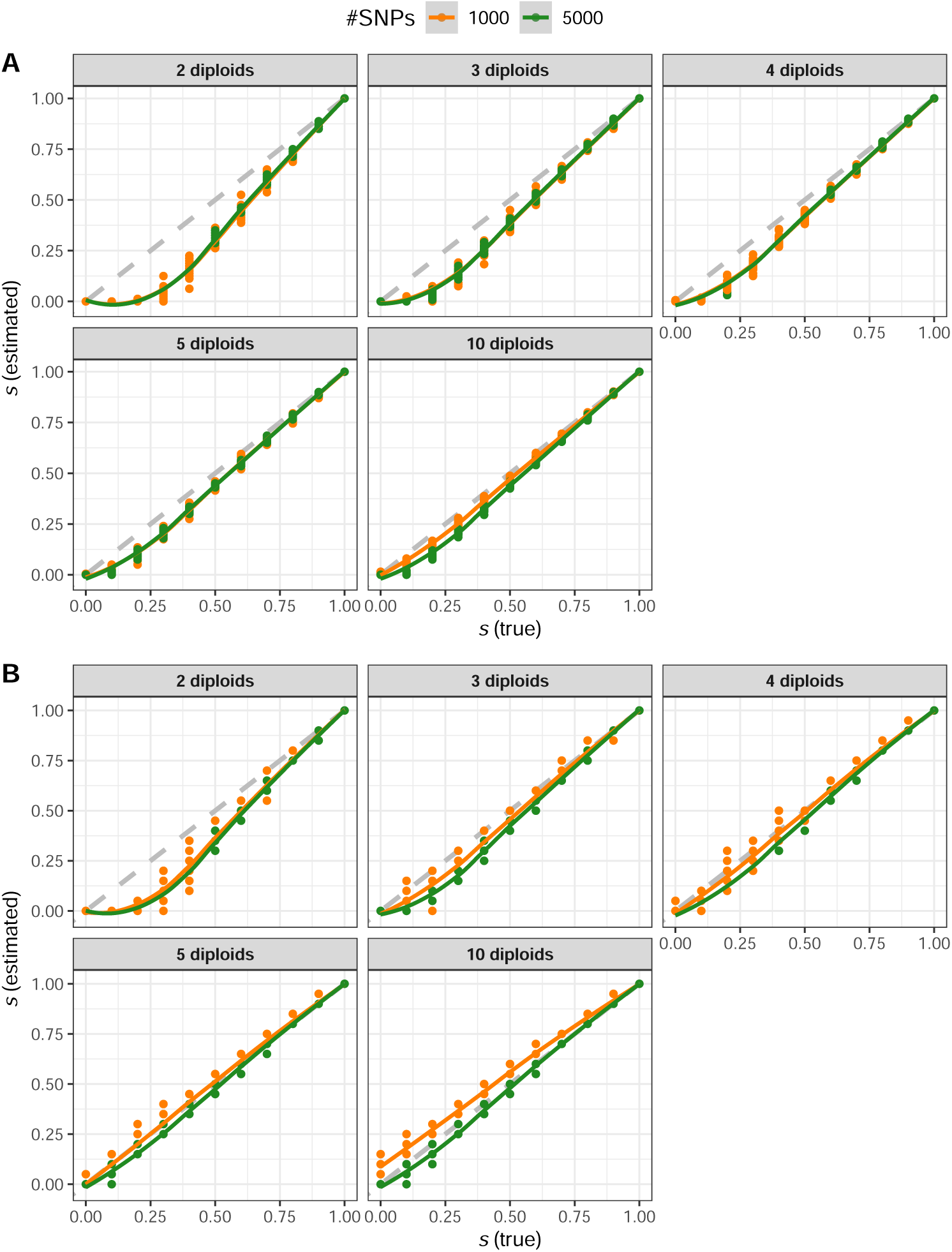
Evaluation of selfing rate (*s*) estimation from simulated genomic data. The x-axis represents the true selfing rate (*s*) used for simulations, while the y-axis shows the selfing rate estimated using our custom method (https://github.com/popgenomics/selfing_ML). **A:** Mean selfing rate estimated across all individuals. **B:** Maximum selfing rate estimated among individuals. Orange and green points represent simulated datasets with 1,000 and 5,000 SNPs, respectively. The number of sampled diploid individuals is indicated in each sub-panel.

**Figure S12:**
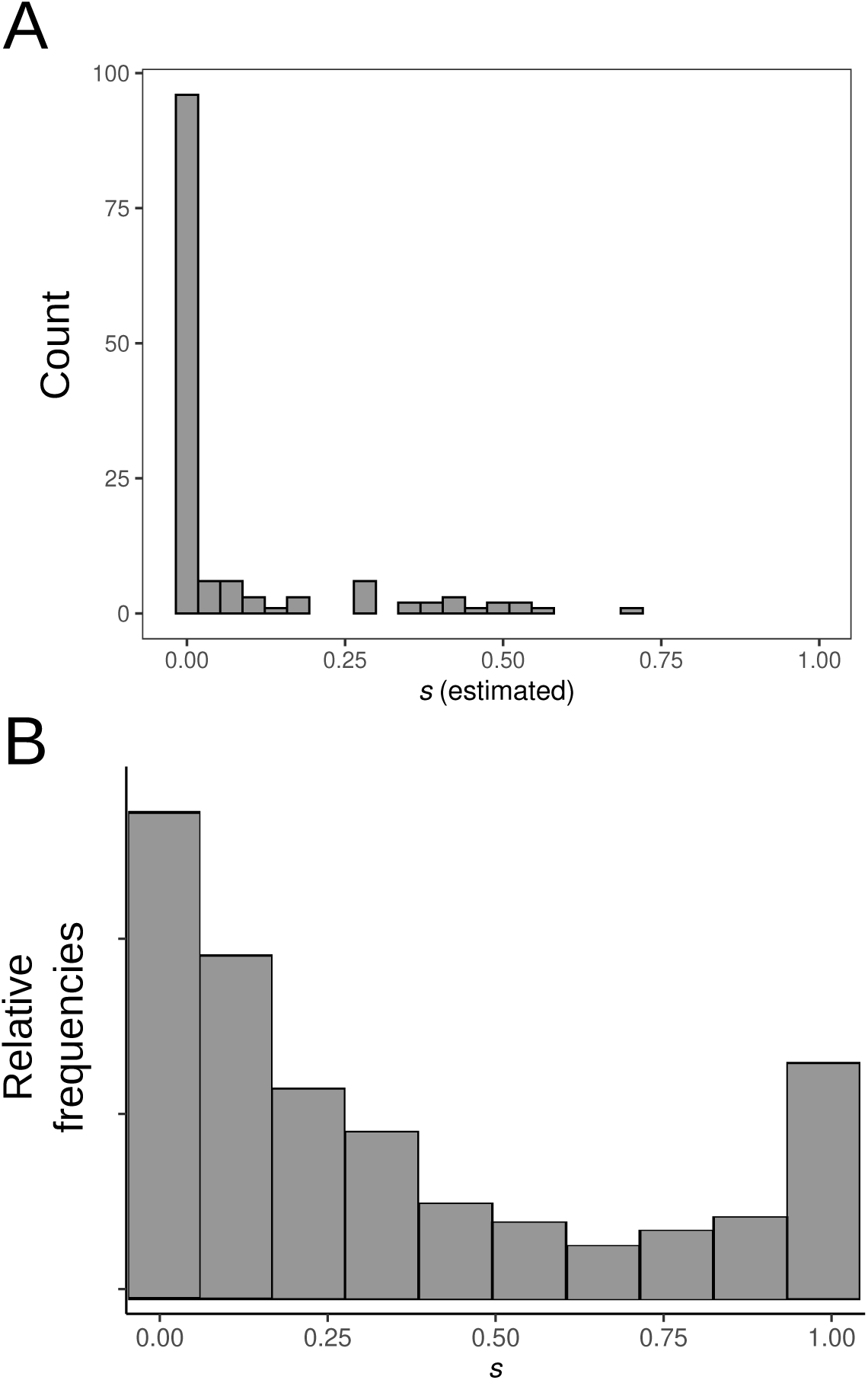
Distribution of selfing rates (*s*). **A:** Distribution of selfing rates among the 135 plant species used in the plant-*versus*-animal comparison, estimated from molecular data. **B:** Meta-analysis of selfing rates across 329 plant species, modified from the study by Igic and Kohn in 2006 (*106*).

**Figure S13:**
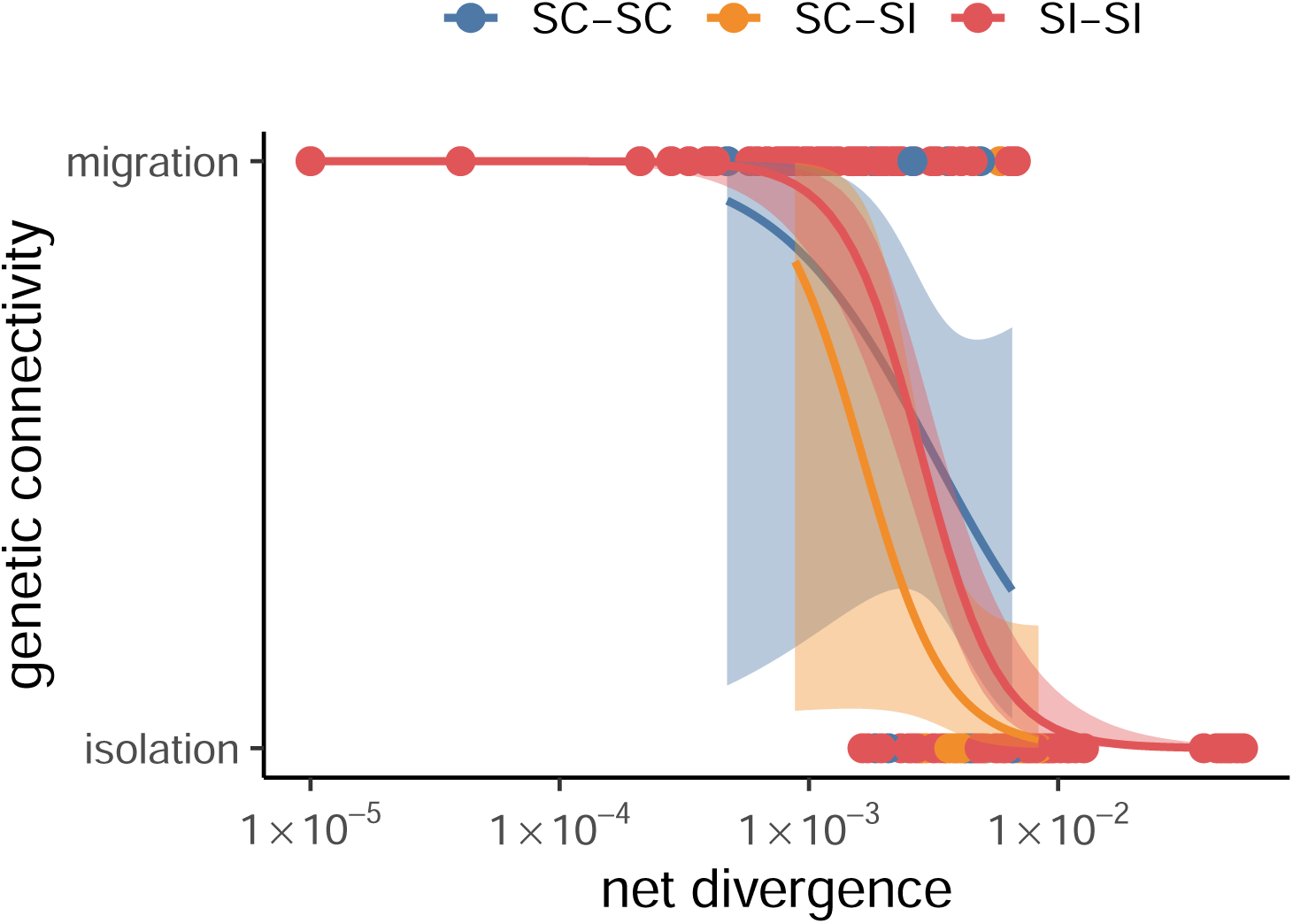
Relationship between mean net divergence and migration/isolation status across plants for different selfing rates. Species pairs were categorized into three groups based on selfing rates: SC-SC (self-compatible pairs), SI-SI (self-incompatible pairs), and SC-SI (mixed pairs). The results of these comparisons are presented in Table S4.

**Figure S14:**
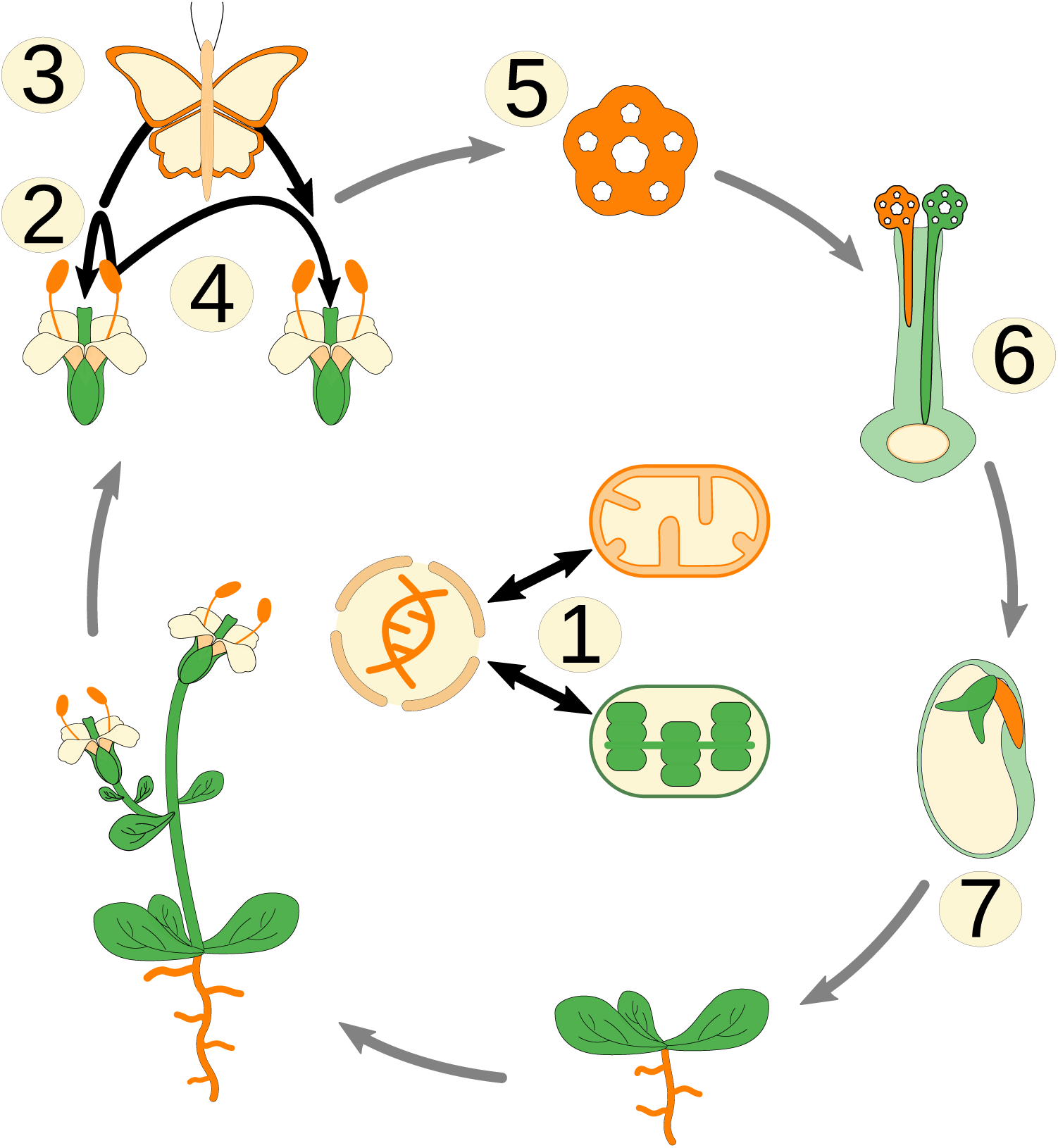
Distinctive attributes of plants and animals that may explain the more rapid accumulation of reproductive isolation in plants. 1. presence of chloroplast and lower cytoplasmic mutation rates than in animals; 2. wide occurrence of self-fertilization in plants; 3. dependence on external pollinators; 4. different dispersion modalities; 5. differences in the strength of haploid selection; 6. predominant exposure to environmental pollen; 7. parental conflicts due to ubiquitous polyandry in plants

### D Supplementary Tables

**Table S1:**
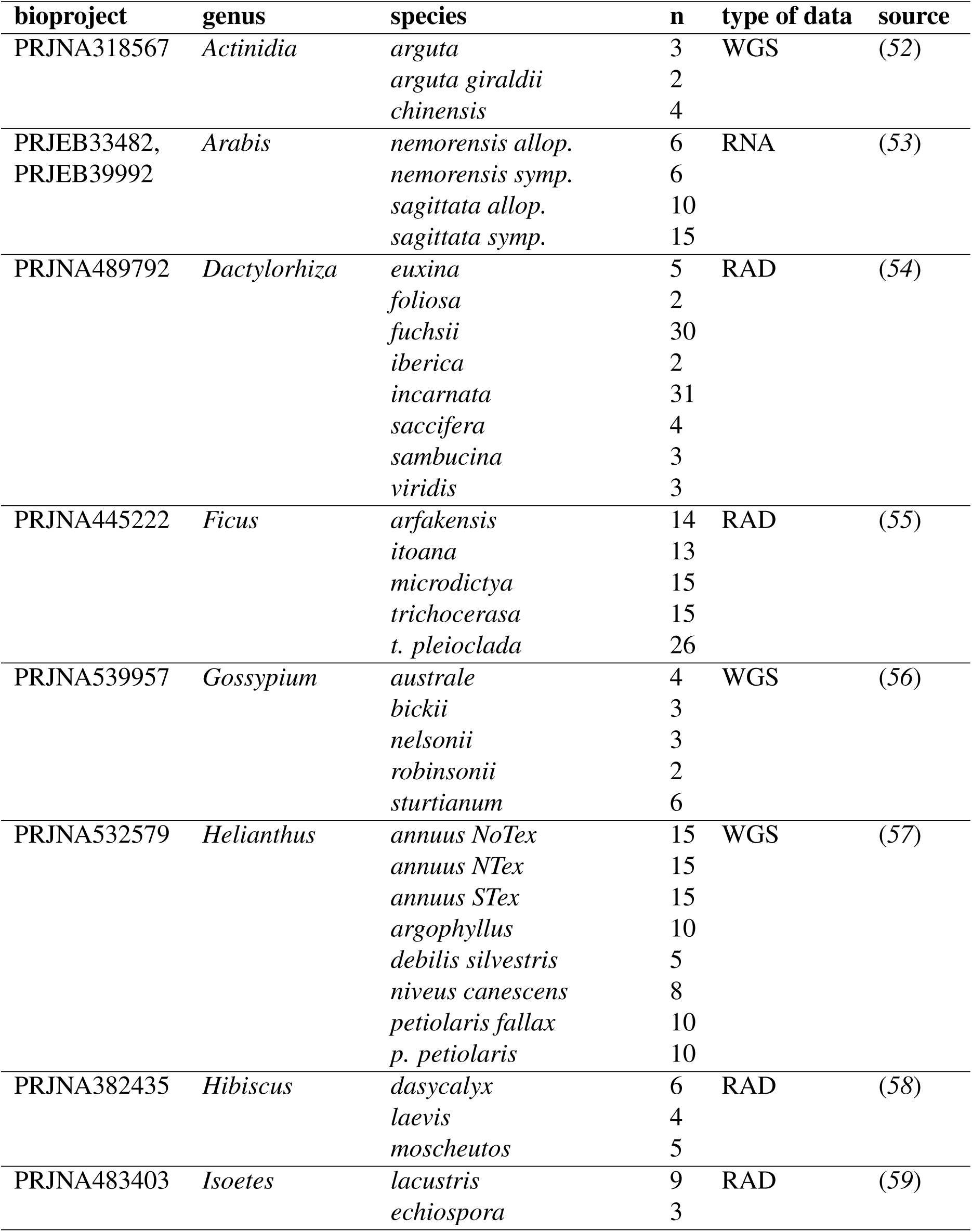
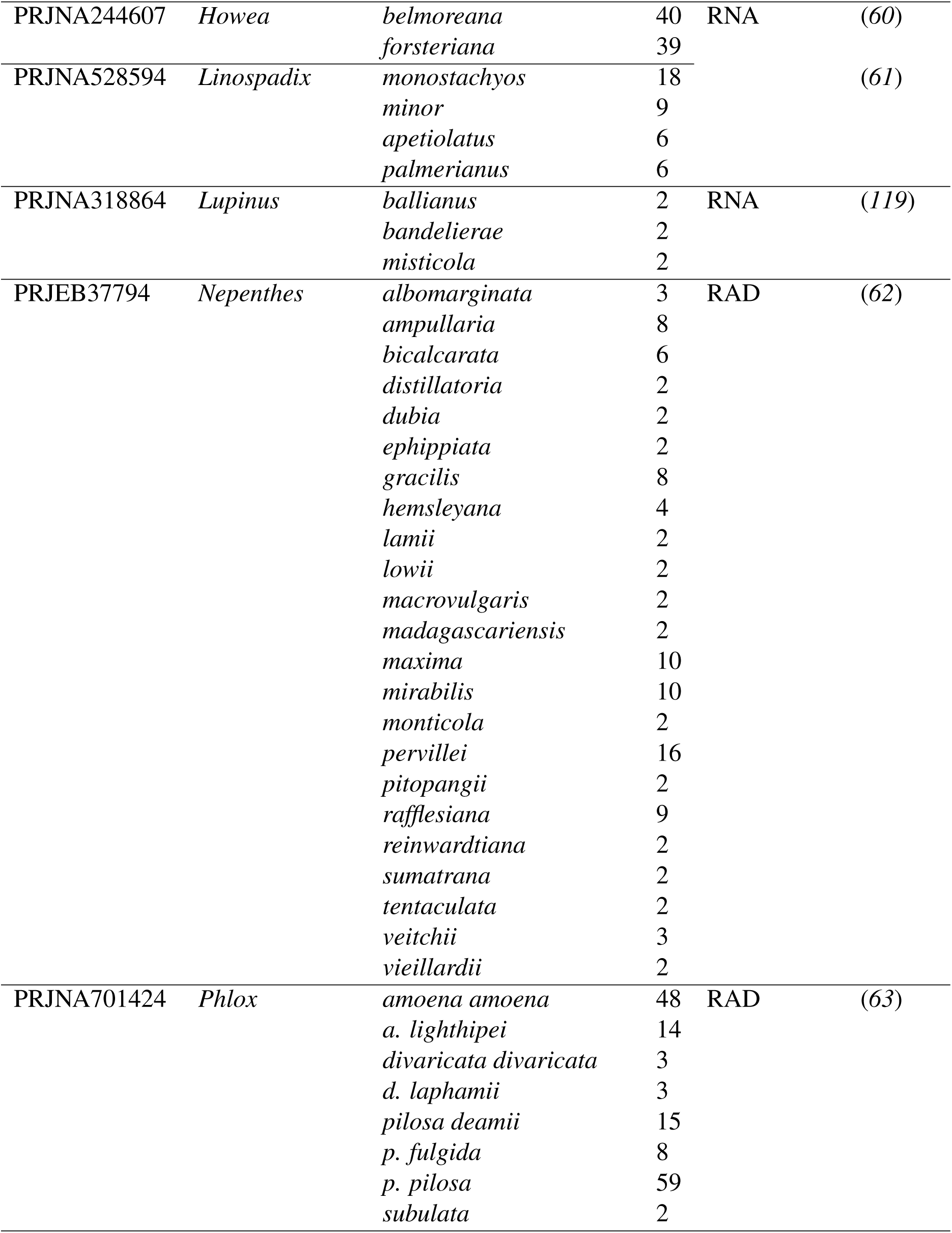
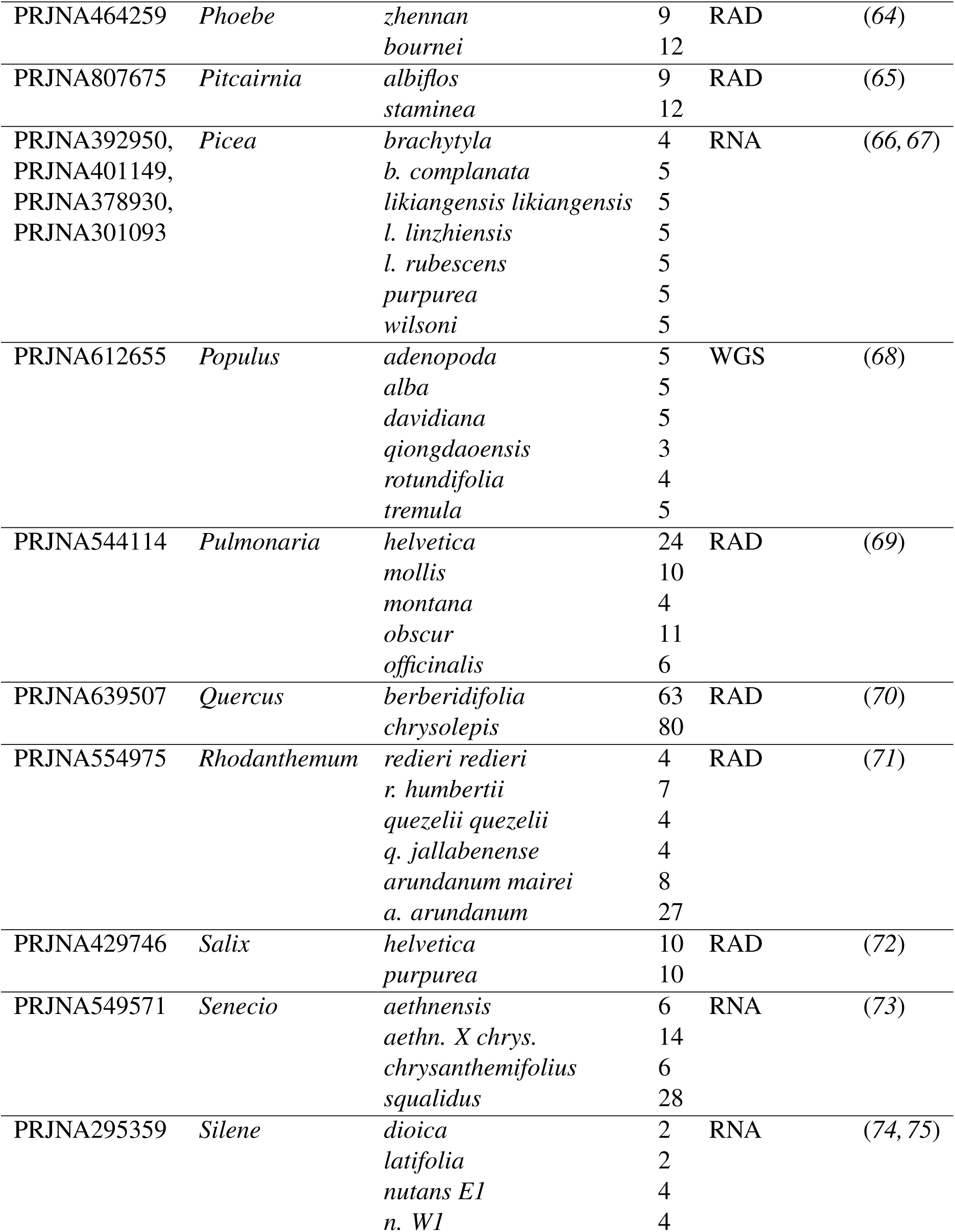
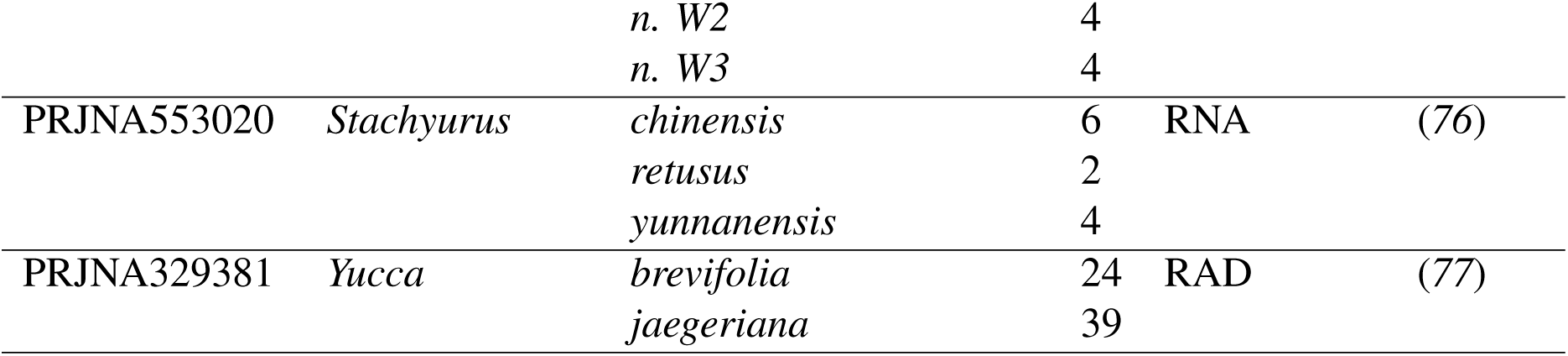
List of retained NCBI datasets.

**Table S2:** Log-likelihood Ratio Test for logit models fitted to plant and animal datasets (Fig.1)

| **model** | $\ell$ | $\beta_0$ | $\beta_1$ | $X_{p=0.5}$ | **P-value** |
| --- | --- | --- | --- | --- | --- |
| $M_0$ | -91.55841 | -11.28733 | -4.50495 | 0.00312 | |
| $M_{plants}$ | -61.60692 | -16.15385 | -6.27316 | 0.00266 | |
| $M_{animals}$ | -8.01012 | -11.13402 | -6.09245 | 0.01488 | |
| | | | | | $< 1 \times 10^{-4}$ |

$\ell$ : log-likelihoods of models $M_0$ , $M_{plants}$ and $M_{animals}$ .
$\beta_0$ : estimated intercept (log10 scaled).
$\beta_1$ : estimated coefficient (log10 scaled).
$X_{p=0.5}$ : inflection point beyond which, for any level of divergence, less than 50% of pairs are expected to be connected by gene flow ( $X_{p=0.5} = 10^{-\frac{\beta_0}{\beta_1}}$ ).
**P-value**: probability that by random chance, the absolute difference between $\ell(M_{plants}) + \ell(M_{animals})$ and $\ell(M_0)$ exceeds the observed value, estimated from 10,000 random permutations.

**Table S3:**
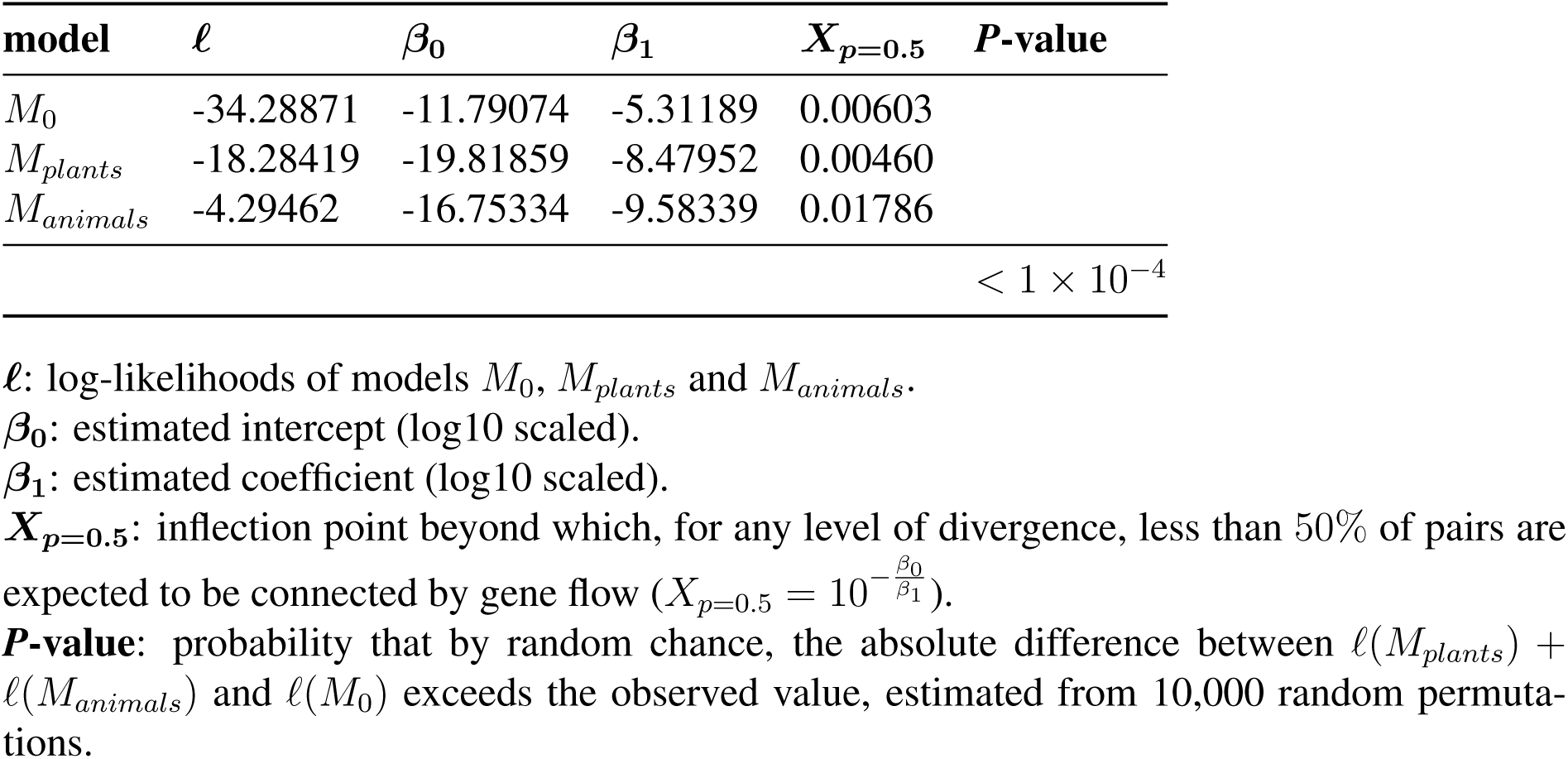
Log-likelihood Ratio Test for logit models fitted to plant and animal datasets obtained by RNA-sequencing only.

**Table S4:**
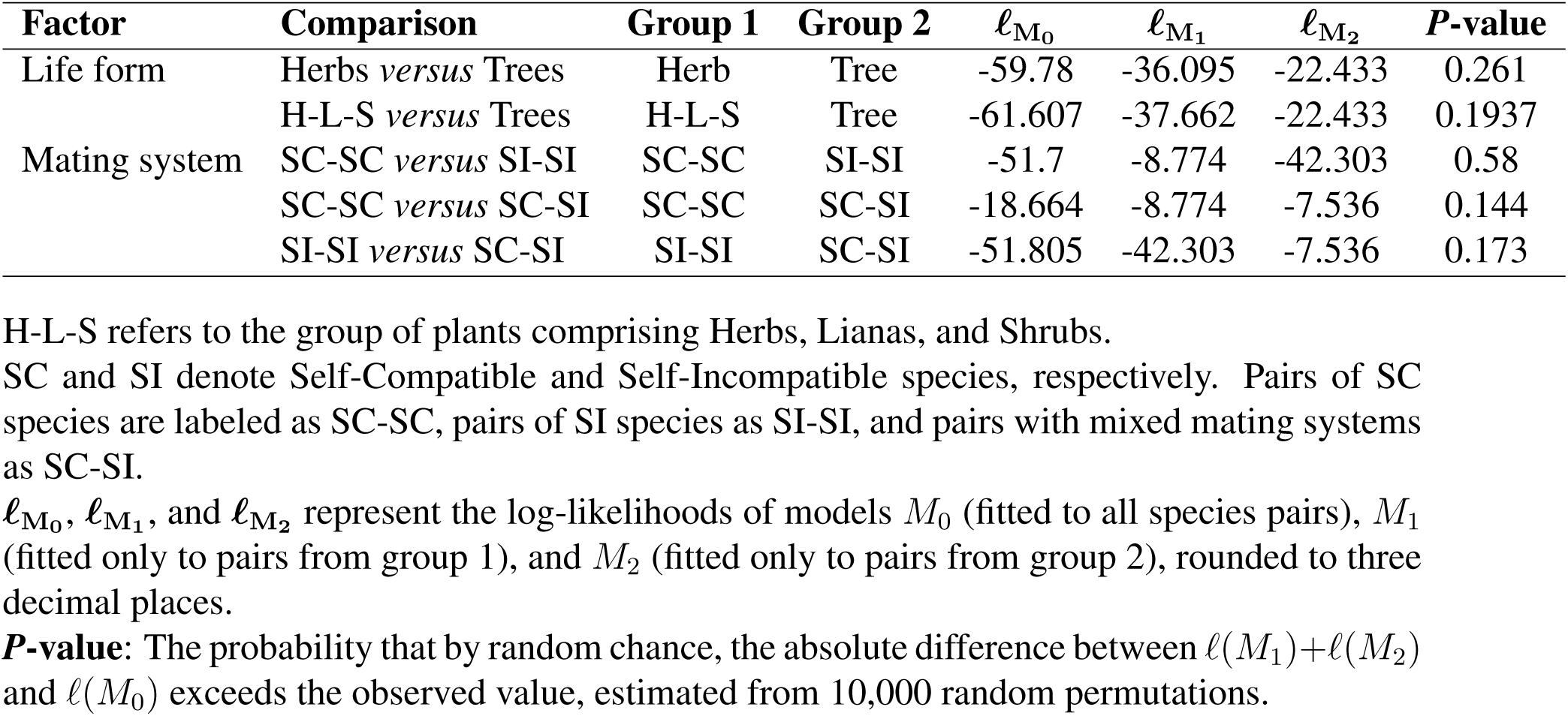
Log-likelihood Ratio Test for logit models assessing factor effects on reproductive isolation dynamics within plants.

## References

1. R. Abbott, et al., Journal of evolutionary biology 26, 229 (2013).

2. G. L. Stebbins, Proceedings of the American Philosophical Society 103, 231 (1959).

3. M. Slatkin, Population genetics and ecology (Elsevier, 1976), pp. 767–780.

4. M. R. Brown, et al., Proceedings of the National Academy of Sciences 120, e2220261120 (2023).

5. A. M. Westram, S. Stankowski, P. Surendranadh, N. Barton, Journal of evolutionary biology 35, 1143 (2022).

6. C.-I. Wu, Journal of evolutionary biology 14, 851 (2001).

7. M. E. Frayer, B. A. Payseur, Evolution 78, 1025 (2024).

8. S. Wright, Annals of eugenics 15, 323 (1949).

9. S. H. Martin, J. W. Davey, C. D. Jiggins, Molecular biology and evolution 32, 244 (2015).

10. A. J. Dagilis, et al., Evolution Letters 6, 344 (2022).

11. C. Fraïsse, et al., Molecular Ecology Resources 21, 2629 (2021).

12. D. R. Laetsch, et al., bioRxiv pp. 2022–10 (2022).

13. V. C. Sousa, M. Carneiro, N. Ferrand, J. Hey, Genetics 194, 211 (2013).

14. C. Roux, et al., PLoS biology 14, e2000234 (2016).

15. C. Fraïsse, et al., Peer Community Journal 2 (2022).

16. E. Mayr, Animal species and evolution (Harvard University Press, 1963).

17. L. Gottlieb, The American Naturalist 123, 681 (1984).

18. P. R. Grant, B. R. Grant, Science 256, 193 (1992).

19. J. Mallet, Trends in ecology & evolution 20, 229 (2005).

20. B. A. Payseur, L. H. Rieseberg, Molecular ecology 25, 2337 (2016).

21. N. Galtier, Evolutionary applications 12, 657 (2019).

22. C. Roux, G. Tsagkogeorga, N. Bierne, N. Galtier, Molecular biology and evolution 30, 1574 (2013).

23. L. C. Moyle, E. B. Graham, Genetics 169, 355 (2005).

24. L. C. Moyle, T. Nakazato, Genetics 179, 1437 (2008).

25. A. L. Sweigart, L. Fishman, J. H. Willis, Genetics 172, 2465 (2006).

26. R. J. Abbott, Journal of Systematics and Evolution 55, 238 (2017).

27. R. G. Harrison, et al., Oxford surveys in evolutionary biology 7, 69 (1990).

28. C. L. Morjan, L. H. Rieseberg, Molecular ecology 13, 1341 (2004).

29. R. Frankham, C. J. Bradshaw, B. W. Brook, Biological Conservation 170, 56 (2014).

30. D. I. Bolnick, et al., Evolution 77, 318 (2023).

31. S. Stankowski, et al., Evolutionary Journal of the Linnean Society 3, kzae001 (2024).

32. S. Greiner, U. Rauwolf, J. Meurer, R. G. Herrmann, Molecular ecology 20, 671 (2011).

33. Z. Postel, et al., Molecular Phylogenetics and Evolution 169, 107436 (2022).

34. A. B. Otten, H. J. Smeets, Human reproduction update 21, 671 (2015).

35. D. Princepe, M. A. de Aguiar, Proceedings of the National Academy of Sciences 121, e2411672121 (2024).

36. C. Goodwillie, S. Kalisz, C. G. Eckert, Annu. Rev. Ecol. Evol. Syst. 36, 47 (2005).

37. E. E. Goldberg, et al., Science 330, 493 (2010).

38. L. Marie-Orleach, C. Brochmann, S. Glémin, PLoS Genetics 18, e1010353 (2022).

39. M. Pickup, et al., New Phytologist 224, 1035 (2019).

40. K. M. Kay, R. D. Sargent, Annual Review of Ecology, Evolution, and Systematics 40, 637 (2009).

41. F. P. Schiestl, P. M. Schlüter, Annual review of entomology 54, 425 (2009).

42. S. Edmands, Molecular ecology 16, 463 (2007).

43. S. Immler, S. P. Otto, The American Naturalist 192, 241 (2018).

44. H. A. Orr, Evolution pp. 1606–1611 (1993).

45. A. Zwaenepoel, H. Sachdeva, C. Fraïsse, Genetics 228, iyae140 (2024).

46. L. Fishman, A. L. Sweigart, Annual review of plant biology 69, 707 (2018).

47. G. D. Sandstedt, A. L. Sweigart, New Phytologist 236, 1545 (2022).

48. M. P. Zuellig, A. L. Sweigart, Evolution 72, 2394 (2018).

49. L. H. Rieseberg, B. K. Blackman, Annals of botany 106, 439 (2010).

50. H. Morlon, et al., Annual Review of Ecology, Evolution, and Systematics 55 (2024).

51. M. François, et al., Rapid establishment of species barriers in plants compared to animals (2024).

52. Y. Liu, et al., New Phytologist 215, 877 (2017).

53. H. Dittberner, A. Tellier, J. de Meaux, Molecular Biology and Evolution 39, msac015 (2022).

54. M. K. Brandrud, et al., Systematic Biology 69, 91 (2020).

55. D. Souto-Vilarós, et al., Journal of Ecology 106, 2256 (2018).

56. C. E. Grover, et al., Genome Biology and Evolution 14, evac170 (2022).

57. G. L. Owens, et al., Molecular Ecology 30, 6229 (2021).

58. J. Norrell (2017).

59. D. P. Wood, J. K. Olofsson, S. W. McKenzie, L. T. Dunning, Botany Letters 165, 476 (2018).

60. L. Dunning, et al., Journal of evolutionary biology 29, 1472 (2016).

61. O. G. Osborne, et al., Molecular Biology and Evolution 36, 2682 (2019).

62. M. Scharmann, A. Wistuba, A. Widmer, Molecular Phylogenetics and Evolution 163, 107214 (2021).

63. B. E. Goulet-Scott, A. G. Garner, R. Hopkins, Evolution 75, 1699 (2021).

64. X. Ding, J. H. Xiao, L. Li, J. G. Conran, J. Li, Journal of Systematics and Evolution 57, 234 (2019).

65. M. M. Tavares, M. Ferro, B. S. S. Leal, C. Palma-Silva, Ecology and Evolution 12, e8834 (2022).

66. Y. Sun, et al., Evolution 72, 2669 (2018).

67. D. Ru, et al., Molecular Ecology 27, 4875 (2018).

68. H. Shang, et al., Philosophical Transactions of the Royal Society B 375, 20190544 (2020).

69. S. Grünig, M. Fischer, C. Parisod, Annals of botany 127, 21 (2021).

70. J. Ortego, L. L. Knowles, Molecular Ecology 29, 4510 (2020).

71. F. Wagner, et al., Molecular phylogenetics and evolution 144, 106702 (2020).

72. S. Gramlich, N. D. Wagner, E. Hörandl, BMC Plant Biology 18, 1 (2018).

73. B. Nevado, S. A. Harris, M. A. Beaumont, S. J. Hiscock, Molecular Ecology 29, 4221 (2020).

74. X.-S. Hu, D. A. Filatov, Molecular ecology 25, 2609 (2016).

75. A. Muyle, et al., Molecular Biology and Evolution 38, 805 (2021).

76. Y. Feng, H. P. Comes, Y.-X. Qiu, Molecular phylogenetics and evolution 150, 106878 (2020).

77. A. M. Royer, M. A. Streisfeld, C. I. Smith, American journal of botany 103, 1730 (2016).

78. B. Langmead, S. L. Salzberg, Nature methods 9, 357 (2012).

79. N. Matasci, et al., Gigascience 3, 2047 (2014).

80. J. M. Catchen, A. Amores, P. Hohenlohe, W. Cresko, J. H. Postlethwait, G3: Genes— genomes— genetics 1, 171 (2011).

81. J. Catchen, P. A. Hohenlohe, S. Bassham, A. Amores, W. A. Cresko, Molecular ecology 22, 3124 (2013).

82. J. R. Paris, J. R. Stevens, J. M. Catchen, Methods in Ecology and Evolution 8, 1360 (2017).

83. L. Excoffier, et al., Bioinformatics 37, 4882 (2021).

84. R. Gutenkunst, R. Hernandez, S. Williamson, C. Bustamante, Nature precedings pp. 1–1 (2010).

85. J. Jouganous, W. Long, A. P. Ragsdale, S. Gravel, Genetics 206, 1549 (2017).

86. N. B. Edelman, et al., Science 366, 594 (2019).

87. M. Malinsky, M. Matschiner, H. Svardal, Molecular ecology resources 21, 584 (2021).

88. T. E. Cruickshank, M. W. Hahn, Molecular ecology 23, 3133 (2014).

89. C. Roux, et al., Journal of Evolutionary Biology 27, 1662 (2014).

90. T. Leroy, et al., New Phytologist 214, 865 (2017).

91. T. Capblancq, et al., bioRxiv pp. 2024–11 (2024).

92. T. Koppetsch, M. Malinsky, M. Matschiner, Systematic biology 73, 769 (2024).

93. J. Ross-Ibarra, et al., PloS one 3, e2411 (2008).

94. R. R. Hudson, Bioinformatics 18, 337 (2002).

95. B. Charlesworth, M. Morgan, D. Charlesworth, Genetics 134, 1289 (1993).

96. N. Barton, B. O. Bengtsson, Heredity 57, 357 (1986).

97. M. A. Beaumont, Annual review of ecology, evolution, and systematics 41, 379 (2010).

98. F. Tajima, Genetics 105, 437 (1983).

99. G. Watterson, Theoretical population biology 7, 256 (1975).

100. F. Tajima, Genetics 123, 585 (1989).

101. M. Nei, W.-H. Li, Proceedings of the National Academy of Sciences 76, 5269 (1979).

102. S. Wright, Genetics 28, 114 (1943).

103. J. Wakeley, J. Hey, Genetics 145, 847 (1997).

104. J. L. Hamrick, J. D. Nason, Forest conservation genetics: Principles and practice pp. 81–90 (2000).

105. F. Austerlitz, S. Mariette, N. Machon, P.-H. Gouyon, B. Godelle, Genetics 154, 1309 (2000).

106. B. Igic, J. R. Kohn, Evolution 60, 1098 (2006).

107. B. Anderson, et al., Iscience 26 (2023).

108. J. Goudet, Molecular ecology notes 5, 184 (2005).

109. A. D. Cutter, New Phytologist 224, 1080 (2019).

110. A. Simon, N. Bierne, J. J. Welch, Evolution Letters 2, 472 (2018).

111. C. Devaux, R. Lande, Journal of evolutionary biology 22, 1460 (2009).

112. R. Tiedemann, et al., Molecular Ecology 13, 1481 (2004).

113. R. Nathan, Science 313, 786 (2006).

114. M. Schilthuizen, M. Giesbers, L. Beukeboom, Heredity 107, 95 (2011).

115. T. Dresselhaus, N. Franklin-Tong, Molecular plant 6, 1018 (2013).

116. R. K. Butlin, C. M. Smadja, The American Naturalist 191, 155 (2018).

117. J. R. Pannell, A.-M. Labouche, Philosophical Transactions of the Royal Society B: Biological Sciences 368, 20120051 (2013).

118. S. Kumar, G. Stecher, M. Suleski, S. B. Hedges, Molecular biology and evolution 34, 1812 (2017).

119. B. Nevado, G. W. Atchison, C. E. Hughes, D. A. Filatov, Nature communications 7, 12384 (2016).

